# MEIS-WNT5A axis regulates development of 4^th^ ventricle choroid plexus

**DOI:** 10.1101/2020.05.07.082370

**Authors:** Karol Kaiser, Ahram Jang, Melody P. Lun, Jan Procházka, Ondrej Machon, Michaela Procházková, Benoit Laurent, Daniel Gyllborg, Renée van Amerongen, Petra Kompaníková, Feizhen Wu, Roger A. Barker, Ivana Uramová, Radislav Sedláček, Zbyněk Kozmík, Ernest Arenas, Maria K. Lehtinen, Vítězslav Bryja

## Abstract

The choroid plexus (ChP) produces cerebrospinal fluid and forms a critical barrier between the brain and the circulation. While the ChP forms in each brain ventricle, it adopts a different shape in each one and remarkably little is known about the mechanisms underlying its development. Here, we show that epithelial WNT5A is critical for determining fourth ventricle (4V) ChP morphogenesis and size. Systemic *Wnt5a* knockout, or forced WNT5A overexpression beginning at E10.5, profoundly reduced the size and development of ChP in all ventricles. However, conditional deletion of *Wnt5a* expression in *Foxj1*-expressing epithelial cells affected only the branched, villous morphology of the 4V ChP. We found that WNT5A was enriched in epithelial cells localized to the distal tips of 4V ChP villi, where WNT5A acted locally to activate non-canonical Wnt signaling via Ror1/Ror2 receptors. During 4V ChP development, MEIS1 bound to the proximal *Wnt5a* promoter, and gain- and loss-of-function approaches demonstrated that MEIS1 regulated *Wnt5a* expression. Collectively, our findings demonstrate a dual function of WNT5A in ChP development and identify MEIS1 and MEIS2 as upstream regulators of *Wnt5a* in the 4V ChP epithelium.

## INTRODUCTION

The choroid plexus (ChP) is a sheet of predominantly epithelial cells that produces cerebrospinal fluid (CSF), secretes factors important for brain development, and forms a critical blood-brain barrier (Chau et al., 2015; Fame and Lehtinen, 2020; Ghersi-Egea et al., 2018; Lehtinen et al., 2011; Silva-Vargas et al., 2016).

ChP tissue is specified during the early stages of brain development (Hunter and Dymecki, 2007), and forms at, or near, the dorsal midline in each ventricle in the brain (lateral ventricle, LV; third ventricle, 3V; fourth ventricle, 4V; Currle et al., 2005). As progenitor cells proliferate and mature into epithelial cells, the epithelial sheet extends in a conveyor belt-like manner into the ventricles (Dani et al., 2019; Liddelow et al., 2010). Interactions between the maturing epithelial cells and the surrounding cellular network of mesenchymal (Wilting and Christ, 1989) and vascular cells (Nielsen & Dymecki, 2010) transform the growing epithelium to adopt its mature form.

The ChP tissues are regionally patterned, and they harbor distinct transcriptomes resulting in ventricle-specific secretomes (Dani et al., 2019; Lun et al., 2015). These findings raise the possibility that local CSF environments are tailored to instruct development of adjacent brain areas. Indeed, the 4V ChP expresses high levels of WNT5A, which is released into the CSF, associates with lipoprotein particles, and influences hindbrain morphogenesis (Kaiser et al., 2019). In mouse, the ChP also appears morphologically distinct in different ventricles; the LV ChP consists of two nearly flat, leaf-like epithelial sheets, while in the 3V and 4V, the ChP adopts a more complex, frond-like structure (Dani et al., 2019). However, despite a general understanding of the key steps required for ChP formation, surprisingly little is known about the molecular mechanisms underlying differences in ChP morphogenesis in these different ventricles.

Wnt5a signaling represents one compelling candidate signaling pathway for regulating ChP development. Systemic *Wnt5a*-deficiency was recently reported to impair ChP development in the LV, 3V and 4V ChP (Langford et al., 2020). However, *Wnt5a* transcription and production is mainly restricted to the epithelium of 4V ChP during development (Kaiser et al., 2019). As such, the molecular mechanisms that regulate the regionalized expression of *Wnt5a* in the ChP and its time- and cell-type restricted functions remain to be elucidated.

Overall, Wnt signaling encompasses highly conserved signaling pathways involved in the regulation of numerous physiological processes during embryonic development and in adulthood (Saito-Diaz et al., 2013). Wnt proteins (WNTs) represent a large family of lipid-modified glycoproteins acting as extracellular ligands that activate either a canonical cascade, mediated by the active β-catenin, or the non-canonical branches of the WNT pathway, depending on ligand binding to a large repertoire of cogent receptors (Niehrs, 2012). Among the WNTs, WNT5A represents a prototypical non-canonical WNT ligand, predominantly linked to the activation of the planar cell polarity (Wnt/PCP) pathway (Kumawat and Gosens, 2015), where it influences various aspects of tissue patterning and establishment of cell polarity (Humphries and Mlodzik, 2018). Notably, WNT5A mediates conserved roles in the outgrowth of body structures including budding tentacles in *Hydra* or limbs and distal digits in mouse (Philipp et al., 2009; Yamaguchi et al., 1999). As such, WNT5A, with its expression mainly restricted to the epithelium of 4V ChP, is an ideal candidate factor for regulating the formation of the complex frond-like shape of the 4V ChP.

Here, we elucidate WNT5A’s dual roles in 4V ChP development. First, at early stages of ChP development, *Wnt5a* temporal expression and dosage are essential for establishing the blueprint for ChP morphogenesis and size. Second, at later stages of ChP development, WNT5A expression and secretion is enriched at the distal tips of 4V ChP epithelium where it acts either locally to activate components of the Wnt/PCP pathway in an autocrine manner or is released into the CSF for long-range signaling. At this later developmental stage, MEIS1 binds to the *Wnt5a* proximal promoter to regulate *Wnt5a* expression in the 4V ChP. Collectively, our findings reveal two distinct developmental stages at which WNT5A plays important roles in establishing 4V ChP form and function.

## RESULTS

### Epithelial WNT5A expression is required for 4V ChP morphogenesis

We previously demonstrated that, during development (E14.5), *Wnt5a* expression is enriched in the mouse 4V ChP epithelium in contrast to the LV ChP (Kaiser et al., 2019) (**Fig. 1A**). *Wnt5a* expression was most prominent at the distal tips of epithelial villi (**Fig. 1B, C**; arrowheads and empty arrowheads). Upon closer inspection of 4V ChP epithelial cells, we observed segregation of WNT5A signal to the apical cell membrane, denoted by Aquaporin-1 (AQP1) staining (**Fig. 1D**, arrowhead), and to the basolateral cell membrane (**Fig. 1D** empty arrowhead), as if poised for local and long-range release. Using a ChP epithelium-based transwell system, we confirmed bi-directional secretion of WNT5A apically and basally by embryonic 4V ChP epithelia (**Supp. Fig. 1A**). Analysis of human fetal brain specimens (week 9 post conception) confirmed WNT5A expression on the apical and basolateral sides of the 4V ChP epithelium (**Fig. 1E**, arrowhead and empty arrowhead, respectively) suggesting *Wnt5a* may have evolutionarily conserved roles in the ChP.

**Figure 1.**
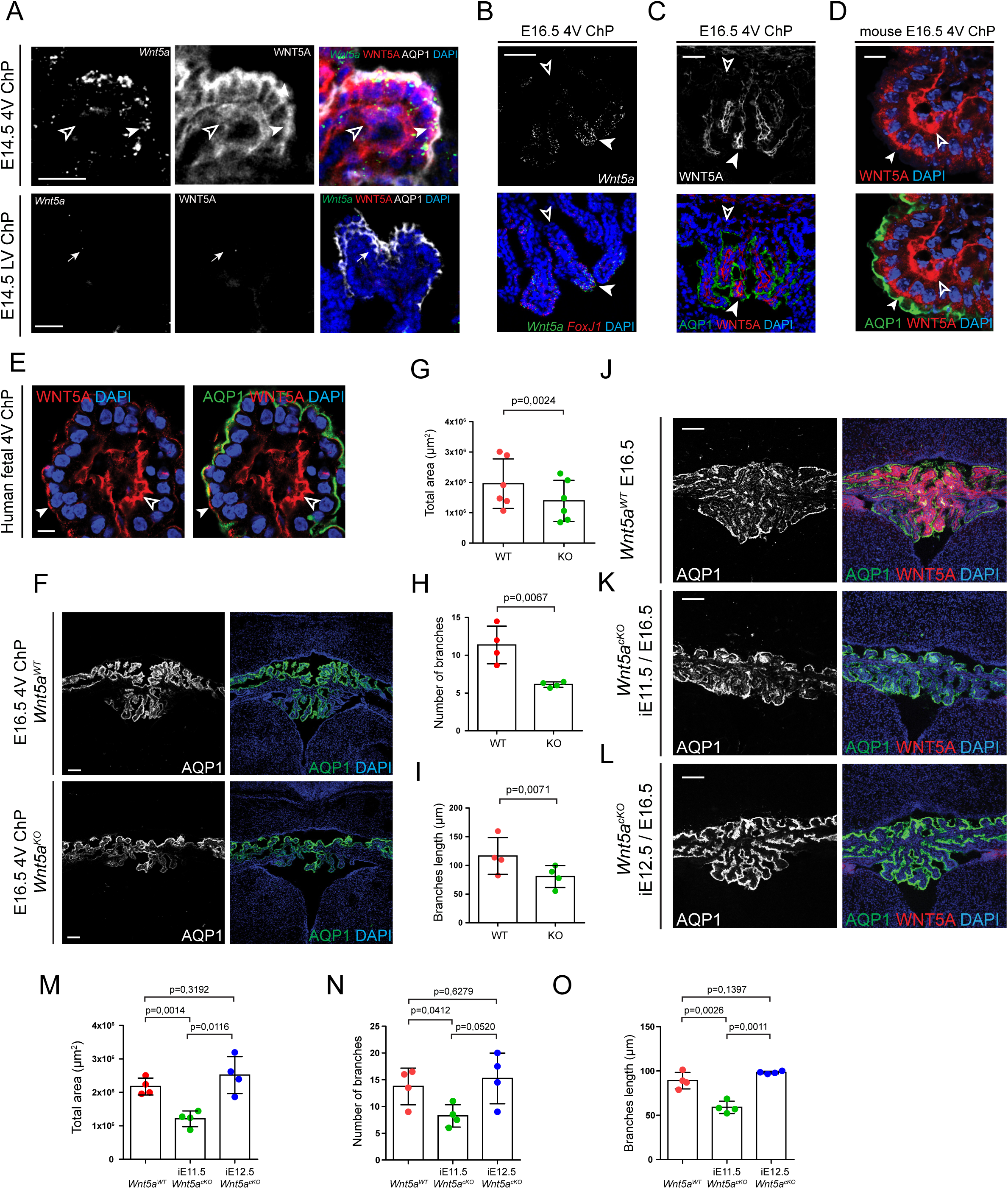
Wnt5a is expressed by 4V ChP epithelium and controls its morphogenesis. (A) *In situ* analysis combined with immunofluorescence staining of E14.5 coronal sections shows *Wnt5a* expression that is restricted only to the ChP epithelium of 4V ChP (arrowhead) with minimal expression in the LV ChP epithelium (arrow). *Wnt5a* expression is largely absent from the stromal compartment (empty arrowhead). Scale bar: 20 μm. (B) *In situ* hybridization analysis of the coronal section showing *Wnt5a* expression restricted to the distal tips (arrowhead) of E16.5 4V ChP epithelium villi as compared to the low *Wnt5a* expression level observed at their base (empty arrowhead). *Foxj1* is a marker of differentiated epithelium. Scale bar: 50 μm. (C) Immunofluorescence analysis reveals that the WNT5A signal is restricted predominantly to the distal tips of E16.5 4V ChP epithelium villi (arrowhead) as opposed to the base of the villi (empty arrowhead). AQP1 is a marker of ChP epithelium. Scale bar: 50 μm. (D, E) Immunofluorescence analysis showing WNT5A signal in both the apical (arrowhead) and basolateral (empty arrowhead) portion of E14.5 mouse (D) and 9-week-old human (E) 4V ChP epithelium. AQP1 is a marker of ChP epithelium. Scale bar: 10 μm. (F) Immunofluorescence analysis of the 4V ChP between *Wnt5a^WT^* vs *Wnt5a^KO^* showing the greatly reduced 4V ChP growth and morphology in *Wnt5a^KO^* embryo at E16.5. Scale bar: 100µm. (G, H, I) Statistical assessment of E16.5 4V ChP morphology indicates there is a significant decrease in total area size (G), the number of branching points (H) and average branch length (I) between *Wnt5a^WT^* vs *Wnt5a^KO^*. Graph shows n=4-6 biologically independent samples; error bars represent mean ± s.d.; P values (two-tailed Student’s t-test with unequal variance). Biological replicates are indicated in the graph. (J, K, L) Immunostaining of E16.5 4V ChP in *Wnt5a^WT^* (J) or *Wnt5a^cKO^* embryos with recombination induced at E11.5 (K) or E12.5 (L). AQP1 staining demonstrates the dramatic reduction in the size and morphology of 4V ChP in *Wnt5a^cKO^* embryos with recombination induced at E11.5 (K) but no effect on branching morphology of 4V ChP in *Wnt5a^cKO^* embryos with recombination induced at E12.5 (L). Scale bar: 100 µm. (M, N, O) Statistical assessment of E16.5 4V ChP morphology indicates there is a significant decrease in total area size (M), the number of branching points (N) and average branch length (O) between *Wnt5a^WT^* vs *Wnt5a^cKO^*. Graph shows n=4 biologically independent samples; error bars represent mean ± s.d.; P values (two-tailed Student’s t-test with unequal variance). Biological replicates are indicated in the graph.

We analyzed the functional consequences of *Wnt5a-*deficiency employing several different mouse models. Phenotypic analysis of *Wnt5a* null mutants (*Wnt5a^KO^*) (Yamaguchi et al., 1999) revealed severe impairment of 4V ChP development as demonstrated by decreased size and reduced branching morphology when compared to the wild-type control embryos (*Wnt5a^WT^*) (**Fig. 1F-I** and **Supp. Fig. 1B**), in agreement with others (Langford et al., 2020). In the most severe *Wnt5a^KO^* cases, the 4V ChP lacked its characteristic convoluted structure and resembled instead the simpler sheet-like shape of the LV ChP (**Supp. Fig. 1C**). We also confirmed the previously reported disruption of LV ChP morphogenesis in *Wnt5a^KO^* embryos which was characterized by collapsed growth and impaired protrusion of the tissue into the lumen of the ventricle (**Supp. Fig. 1D**) accompanied by general shortening of the tissue (**Supp. Fig. 1E, F**). We investigated whether the observed morphological differences in 4V ChP were due to altered proliferative or apoptotic activity. No differences were observed in the EdU incorporation rate (delivered at E13.5 and analyzed at E16.5) when markers of proliferation (KI67) in mature epithelial cells (AQP1) were examined in the 4V ChP epithelium of *Wnt5a^KO^* ChP (**Supp. Fig. 1 G-I**). Similarly, no changes in apoptotic activity analyzed by cleaved caspase 3 (CASP3) staining were detected in the embryonic 4V ChP between *Wnt5a^KO^* and control littermates (**Supp. Fig. 1J**), suggesting that any contributions in these processes to the phenotypes observed must have occurred earlier in development.

Next, we conditionally knocked out *Wnt5a* (*Wnt5a^cKO^*) in ChP epithelial cells during earlier stages of ChP development using an inducible *Foxj1* promoter-driven CRE^ERT2^ system (Kaiser et al., 2019) (**Supp. Fig. 2A, B**). In line with the reported properties of the Cre-lox recombination system (Nakamura et al., 2006) we observed the loss of WNT5A protein approximately 24 hours following tamoxifen injection **(Supp. Fig. 2C)**. Tissue-specific ablation of WNT5A in 4V ChP epithelium of *Wnt5a^cKO^* vs. control embryos persisted at least till E16.5 when tamoxifen is injected at E12.5 (**Supp. Fig. 2D)**. WNT5A was readily detected in the domain adjacent to the 4V ChP where *FoxJ1* is not expressed in both E16.5 *Wnt5a^WT^* and *Wnt5a^cKO^* embryos (**Supp. Fig. 2E**). The concomitant loss of WNT5A immunoreactivity in the stromal compartment of the *Wnt5a^cKO^* 4V ChP strongly suggested ChP epithelial cells as the primary source of WNT5A for this compartment (**Supp. Fig. 2E**).

**Figure 2.**
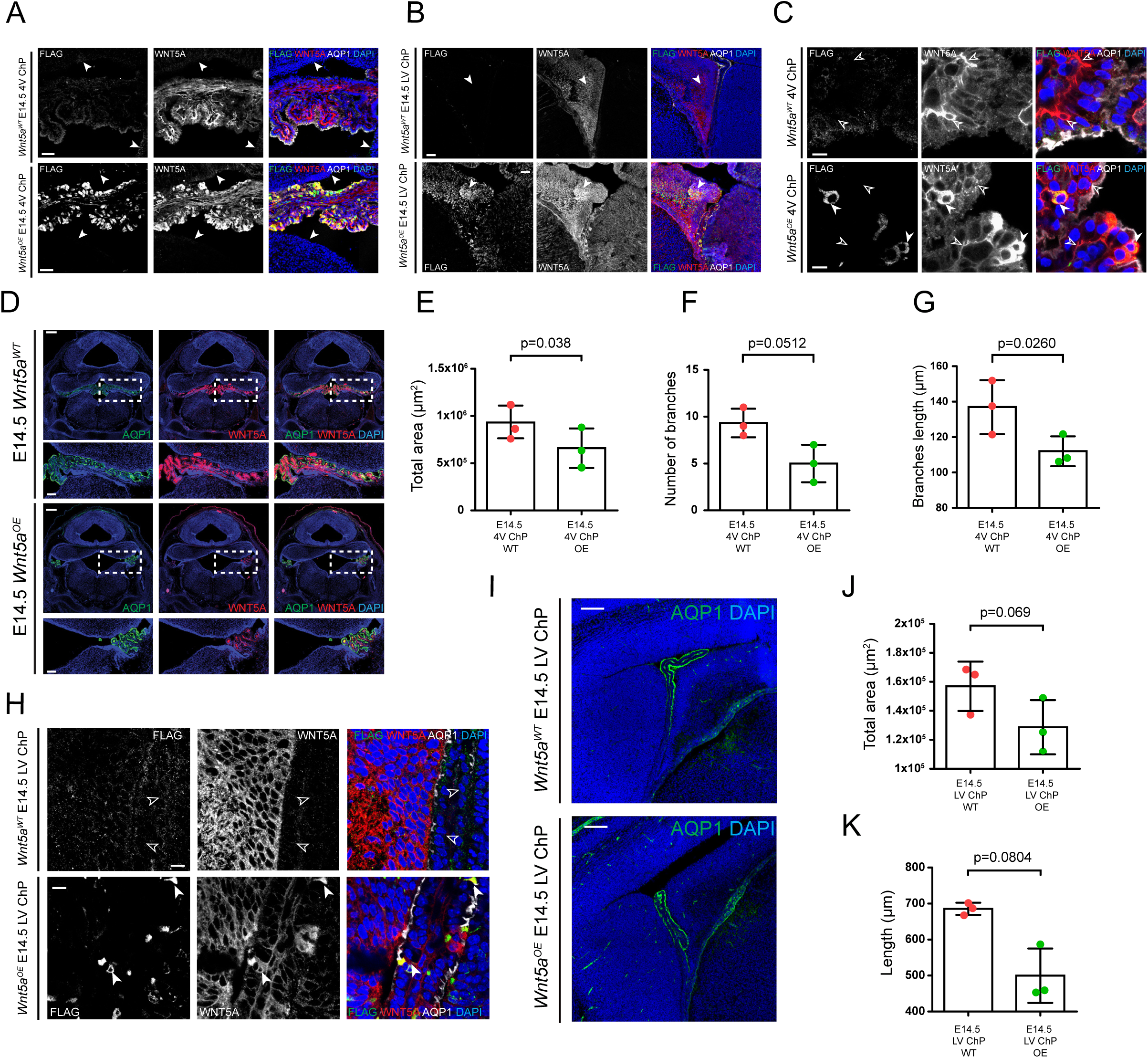
Ectopic Wnt5a expression impairs morphogenesis of lateral and 4V ChP. (A) Immunofluorescence analysis of coronal sections showing an induction of Flag-tagged WNT5A (FLAG-WNT5A) overexpression in 4V ChP in *Wnt5a^OE^* embryos as compared to *Wnt5a^WT^* littermates. AQP1 is a marker of ChP epithelium. Scale bar: 50 μm. (B) Immunofluorescence analysis of coronal sections showing induction of FLAG-WNT5A overexpression in LV ChP in Wnt5a*^OE^* embryos as compared to *Wnt5a^WT^* littermates. Strong induction of ectopic FLAG-WNT5A protein can be also observed in the adjacent Cortical Hem (CH; arrowhead). AQP1 is a marker of ChP epithelium. Scale bar: 50 μm. (C) Higher magnification view showing that the signal for the ectopically induced FLAG-WNT5A is mostly restricted to the cytoplasm of cells producing FLAG-WNT5A in 4V ChP epithelium (arrowheads), while no FLAG-WNT5A can be detected in the surrounding basal lamina or stroma as compared to endogenous WNT5A (empty arrowheads). AQP1 is a marker of ChP epithelium. Scale bar: 10 μm. (D) Immunofluorescence analysis of coronal sections demonstrates the reduced 4V ChP morphology between *Wnt5a^WT^* vs Wnt5a*^OE^* at E14.5. Inset displays higher magnification view of the 4V ChP (dotted box) in both *Wnt5a^WT^* and Wnt5a*^OE^* embryos. AQP1 is a marker of ChP epithelium. Scale bar: 300 µm. Panel scale: 100 µm. (E, F, G) Statistical assessment of E14.5 4V ChP morphology indicates that there is a significant decrease in total area size (E), the number of branching points (F) and average branch length (G) between *Wnt5a^WT^* vs Wnt5a*^OE^*. Graph shows n=3 biologically independent samples; error bars represent mean ± s.d.; P values (two-tailed Student’s t-test with unequal variance). Biological replicates are indicated in the graph. (H) Higher magnification view shows that ectopic production of FLAG-WNT5A in Wnt5a*^OE^* embryos results in detection of the WNT5A signal in LV ChP epithelial cells (arrowheads) that normally display very low levels of WNT5A signal as observed in *Wnt5a^WT^* LV ChP epithelium (empty arrowheads). AQP1 is a marker of ChP epithelium. Scale bar: 10 μm. (I) Immunofluorescence analysis of coronal sections demonstrates altered LV ChP morphology between *Wnt5a^WT^* vs Wnt5a*^OE^* at E14.5. AQP1 is a marker of ChP epithelium. Scale bar: 100 µm. (J, K) Statistical assessment of E14.5 LV ChP morphology indicates that there is a less robust decrease (as observed for 4V ChP) in total area size (J) and average length (K) between *Wnt5a^WT^* vs Wnt5a*^OE^*. Graph shows n=3 biologically independent samples; error bars represent mean ± s.d.; P values (two-tailed Student’s t-test with unequal variance). Biological replicates are indicated in the graph.

The specific timing of *Wnt5a* ablation had strikingly different effects on the developing 4V ChP. Induction of Cre recombination by tamoxifen injection at E11.5 profoundly disrupted the overall size and branching morphology of the 4V ChP in *Wnt5a^cKO^* (**Fig. 1J, K**), when analyzed at E16.5. In contrast, induction of Cre recombination at E12.5 failed to induce a visible phenotype in the E16.5 4V ChP (**Fig. 1L-O**). The morphological defects observed following E11.5 Cre-recombination were regionalized within the 4V ChP, being most pronounced in the ventral part of the 4V ChP, located next to the lower rhombic lip (LRL) region as compared to the more lateral region of the ChP adjacent to the upper rhombic lip (URL) region (**Supp. Fig. 2F**, arrowheads). Consistent with the systemic *Wnt5a^KO^*-embryos, no obvious differences in either proliferative (EdU+ and KI67+ cells) or apoptotic cell death (CASP3) markers were again observed at E14.5 (**Supp. Fig. 2G, H**). In contrast with the findings in the systemic *Wnt5a^KO^* embryos (**Supp. Fig. 1 D-F**), we did not detect any morphological changes in the developing LV ChP in *Wnt5a^cKO^* embryos (**Supp. Fig. 2I-K**). Taken together, these data demonstrate that the *Wnt5a* expression between E11.5 and E12.5 is essential for establishing the blueprint for 4V ChP morphogenesis and size. In turn, *Wnt5a*-deficiency at this early stage has profound, long-lasting consequences on ChP morphology, and as such, differences in proliferation and apoptotic cell death were not apparent when examined at later ages.

### Wnt5a overexpression during late embryogenesis results in defective morphogenesis of embryonic ChPs

*Wnt5a* dosage, demonstrated by both ablation and ectopic overexpression studies, was previously reported to affect the morphogenesis of the small intestine (Cervantes et al., 2009, van Amerongen et al., 2012), a simple columnar epithelium with similarities to the ChP (Grosse et al., 2011). To determine the consequences of supplemental WNT5A on the 4V ChP, we employed a previously established model for *Wnt5a* overexpression (van Amerongen et al., 2012) (*Wnt5a^OE^*). In this system, global overexpression is driven by the *Rosa26* promoter. In these mice, ectopic FLAG-WNT5A expression matched the spatial pattern of endogenous WNT5A distribution we had observed under normal physiological conditions (**Fig. 1J** and **Supp. Fig. 2D**)(Kaiser et al., 2019). Higher FLAG-WNT5A was detected in embryonic 4V ChP vs. surrounding regions (**Fig. 2A**; arrowheads). We also observed FLAG-WNT5A expression in other brain regions that typically express *Wnt5a* including the cortical hem (**Fig.2B** and **Supp. Fig. 3A**; arrowheads). In the 4V ChP, FLAG-WNT5A localized to the cytoplasm of epithelial cells and revealed limited local spreading within the surrounding 4V ChP extracellular space (**Fig. 2C**, arrowheads and empty arrowheads).

**Figure 3.**
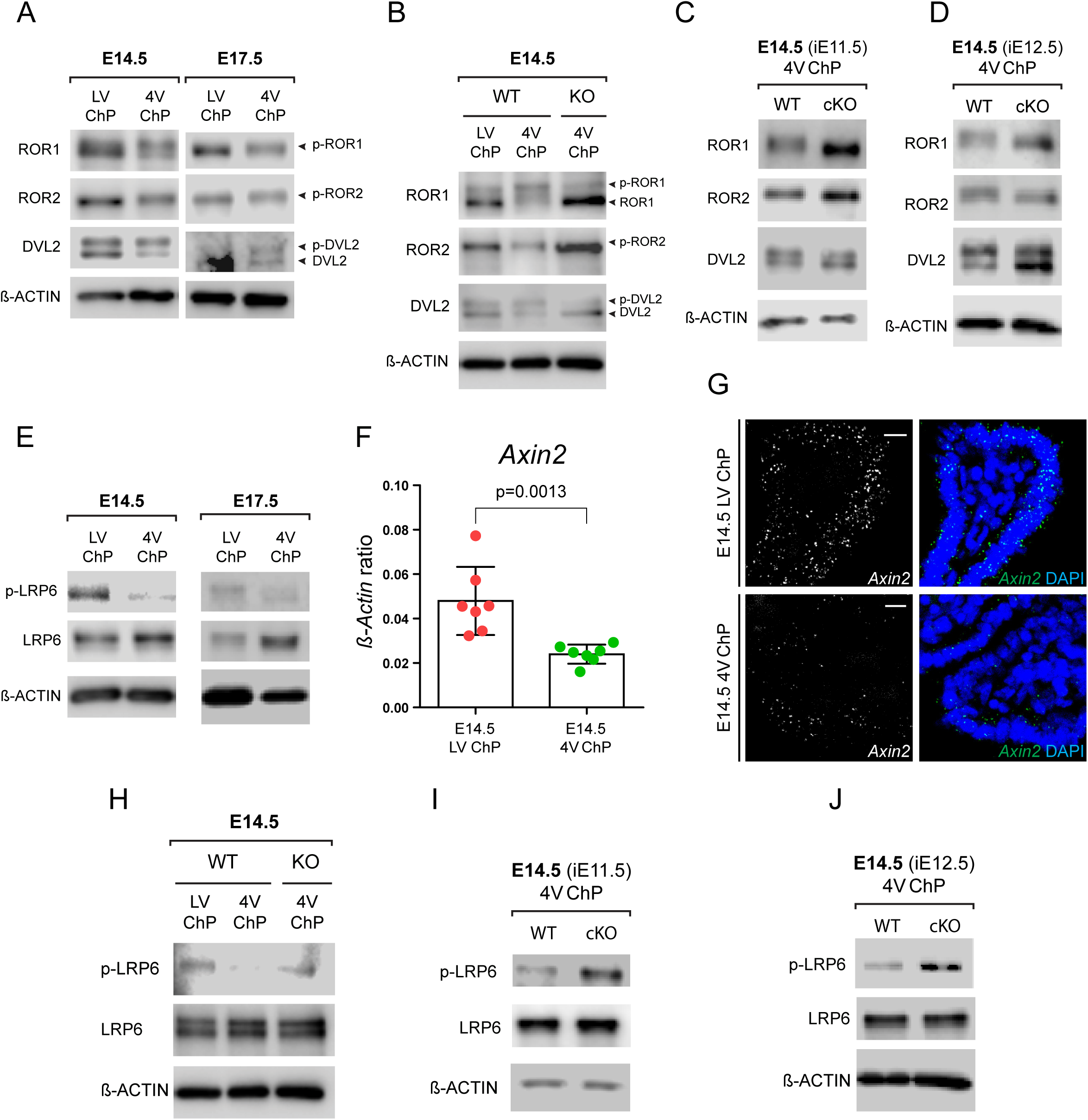
WNT5A activates non-canonical Wnt signaling in 4V ChP. (A) Western blot analysis of LV ChP and 4V ChP at E14.5 and E17.5 displays persistent pattern of non-canonical Wnt signaling pathway induction specific for 4V ChP as compared to the LV ChP. Phosphorylation-dependent shift in the readouts of WNT5A-triggered signaling (ROR1, ROR2 and DVL2) is indicated by the arrowheads. β-actin serves as a loading control. (B) Western blot analysis of E14.5 LV ChP and 4V ChP isolated from *Wnt5a^WT^* and *Wnt5a^KO^* embryos indicates loss of non-canonical Wnt signaling component (ROR1, ROR2 and DVL2) activation upon ablation of WNT5A. Phosphorylation-dependent shift in the readouts of WNT5A-triggered signaling is indicated by the arrowheads. β-actin serves as a loading control. (C, D) Western blot analysis of *Wnt5a^WT^* and either E11.5-induced (C) or E12.5-induced (D) *Wnt5a^cKO^* embryos highlights absence of non-canonical Wnt pathway activation upon loss of WNT5A-mediated signaling, specifically in 4V ChP epithelium. β-actin serves as a loading control. (E) Western blot analysis of LV ChP and 4V ChP at E14.5 and E17.5 shows LV ChP-specific activation of the canonical Wnt signaling pathway as demonstrated by the increase of phosphorylated LRP6 (pLRP6) protein level. β-actin serves as a loading control. (F) Real-time qPCR of Wnt canonical target gene *Axin2* expression in LV ChP and 4V ChP at E14.5. The expression was normalized against expression levels of *β-actin* in each condition. Graph shows n=7 biologically independent samples; error bars represent mean ± s.d.; P values (two-tailed Student’s t-test with unequal variance). Biological replicates are indicated in the graph. (G) *In situ* hybridization analysis of the coronal sections confirming increased *Axin2* expression in E14.5 LV ChP epithelium as compared to 4V ChP epithelium. Scale bar: 10 μm. (H) Western blot analysis of E14.5 LV ChP and 4V ChP isolated from *Wnt5a^WT^* and *Wnt5a^KO^* embryos highlights partial activation of canonical Wnt signaling in *Wnt5a^KO^* 4V ChP as measured by the increase in pLRP6 protein level. β-actin serves as a loading control. (I, J) Western blot analysis of *Wnt5a^WT^* and either E11.5-induced (I) or E12.5-induced (J) *Wnt5a^cKO^* embryos shows minor activation of canonical Wnt signaling upon 4V ChP epithelium specific ablation of WNT5A as indicated by the increase of overall pLRP6 protein levels.

*Wnt5a* overexpression had detrimental consequences on the formation of the 4V ChP that were characterized by considerable reduction of its overall size and decreased complexity of its branched, villous structure (**Fig. 2D-G**). This effect probably relates to overactivation of WNT5A signaling early on in epithelial cells where future 4V ChP will arise as evidenced by presence of AQP1+ cells (**Supp. Fig. 3B**). As with the knockout experiments, we did not observe any changes in Ki67 or CASP3 immunoreactivity in the E14.5 4V ChP of *Wnt5a^OE^* embryos as compared to *Wnt5a^WT^* littermate controls (**Supp. Fig. 3C, D**, arrowheads). We also noted increased FLAG-WNT5A expression in LV ChP epithelial cells of *Wnt5a^OE^* embryos as compared to controls (**Fig. 2H**, arrowheads and empty arrowheads). Only minor effects of *Wnt5a* overexpression on LV ChP size and length were observed (**Fig. 2I-K**). Taken together, our data demonstrate that ChP morphogenesis relies on tightly regulated WNT5A-dosage, likely stipulated at the earliest stages of ChP development.

### WNT5A promotes activation of non-canonical WNT signaling in 4V ChP

At E16.5, WNT5A localizes to the distal tips of the 4V ChP epithelium (**Fig. 1B, C**) and can be released apically into the CSF for long-range signaling to instruct hindbrain development (Kaiser et al., 2019). In addition, WNT5A can be released basally (**Fig.1 D-E** and **Supp. Fig. 1A**) for potential local signaling within the ChP. To address the extent of WNT5A signaling activation we performed biochemical analyses of LV and 4V ChP tissue lysates. The activation leads to phosphorylation of several downstream signaling pathway components - namely WNT5A receptors ROR1 and ROR2 (Grumolato et al., 2010; Ho et al., 2012; Yamamoto et al., 2007) and a key signaling mediator Dishevelled-2 (DVL2) (Bryja et al., 2007).This activation can be detected by Western blotting as a phosphorylation-dependent mobility shift. Using this approach, we discovered considerably strong activation of Wnt signaling in the embryonic 4V ChP compared to LV ChP during late embryogenesis (**Fig. 3A** and **Supp. Fig. 4A**). Knock out of WNT5A in HEK293 cells mimicked the differences between 4V and LV ChP suggesting that activation of these readouts depended on WNT5A (**Supp. Fig. 4B**). Indeed, *Wnt5a* ablation in both *Wnt5a^KO^* and *Wnt5a^cKO^* ChPs resulted in the loss of induction of non-canonical WNT pathway components, thus demonstrating that their activation was mediated exclusively by 4V ChP epithelium derived WNT5A (**Fig. 3B-D**). Conversely, we observed that LV ChP, which is normally devoid of endogenous WNT5A-signalling (Kaiser et al., 2019), exhibited increased activation of various markers of canonical Wnt signaling including upregulation of downstream target genes *Axin2* (Jho et al., 2002) and *Lef1* (Hovanes et al., 2001) or phosphorylation of LRP6 (Tamai et al., 2000) as compared to 4V ChP (**Fig. 3E-G** and **Supp. Fig. 4C-E**). We attribute these effects to the capacity of WNT5A to suppress the canonical WNT pathway during development (Topol et al., 2003). In support of this model, ablation of *Wnt5a* using both *Wnt5a^KO^* and *Wnt5a^cKO^* mouse models resulted in a partial shift in balance of WNT signaling from non-canonical to canonical signaling (**Fig. 3H-J**). The progressive decrease of canonical Wnt signaling target genes *Axin2* and *Lef1* expression along the proximal-distal (P-D) axis was correlated with gradually increasing expression levels of *Wnt5a* within 4V ChP epithelium (**Supp. Fig. 4F**).

**Figure 4.**
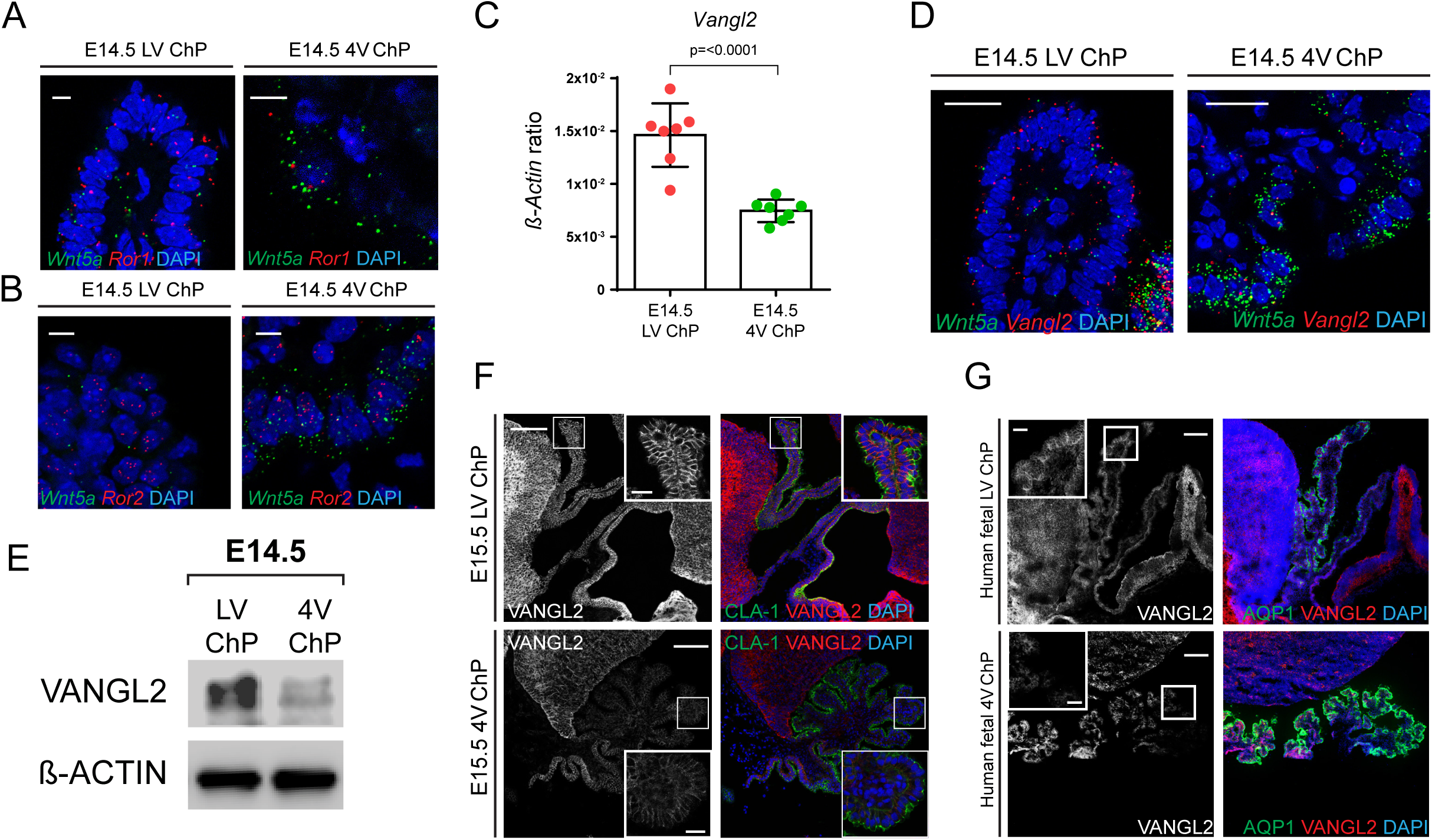
Wnt/PCP components are expressed in embryonic ChP epithelium. (A, B) *In situ* hybridization analysis of the coronal sections showing gene expression of downstream non-canonical Wnt pathway components *Ror1* (A) and *Ror2* (B) that is restricted to both E14.5 LV and 4V ChP epithelium. *Wnt5a* expression highlights ChP epithelium. Scale bar: 5 μm. (C) Real-time qPCR of *Vangl2* expression in LV ChP and 4V ChP at E14.5. The expression was normalized against expression level of *β-actin* in each condition. Graph shows n=7 biologically independent samples; error bars represent mean ± s.d.; P values (two-tailed Student’s t-test with unequal variance). Biological replicates are indicated in the graph. (D) *In situ* hybridization analysis of the coronal sections showing higher gene expression level of *Vangl2* at E14.5 in LV ChP compared to 4V ChP epithelium. *Wnt5a* expression highlights ChP epithelium. Scale bar: 20 μm. (E) Western blot analysis of VANGL2 protein levels between LV ChP and 4V ChP, isolated at E14.5, showing higher VANGL2 protein abundance in the LV ChP as compared to the 4V ChP at E14.5. β-actin serves as a loading control. (F) Immunofluorescence analysis of sagittal sections demonstrating differential VANGL2 protein levels between E14.5 LV ChP and 4V ChP epithelium with increased signal intensity observed in LV ChP epithelium, especially in its most distal portion (Inset). VANGL2 signal segregates predominantly to the apical portion of the basolateral compartment at the border of neighboring epithelial cells (Insets). Claudin-1 (CLA-1) is a marker of ChP epithelium. Scale bar: 200 μm. Inset scale bar: 20 μm. (G) Immunofluorescence analysis of sagittal sections demonstrates differential VANGL2 protein levels between 9 week old human fetal LV ChP and 4V ChP epithelium with increased signal intensity observed in LV ChP epithelium, especially in its most distal portion (Inset). AQP-1 is a marker of ChP epithelium Scale bar: 200 μm. Inset scale bar: 50 μm.

### Embryonic ChP epithelium expresses various WNT/PCP components

We analyzed in greater detail the spatial distribution of Wnt pathway signaling components in the ChP. Expression of genes encoding non-canonical WNT pathway components, including *Dvl2* and both *Ror1* and *Ror2*, were restricted to the epithelial cell layer and not detected in the stromal compartment of the LV and 4V ChP (**Fig. 4A, B** and **Supp. Fig. 5A-B’**). Given WNT5A’s central role in the WNT/PCP pathway, we also inspected the expression pattern of *Vangl2*, a downstream mediator of WNT5A dependent signaling in the WNT/PCP pathway (Gao et al., 2011). *Vangl2* expression exhibited a distinct spatial pattern with higher levels detected in the embryonic LV ChP compared to the 4V ChP (**Fig. 4C**). In addition, *Vangl2* expression was restricted to ChP epithelial cells and was absent from the stroma, in agreement with our observations regarding expression of genes encoding other non-canonical Wnt components (**Fig. 4D** and **Supp. Fig. 5B’’**). We validated a VANGL2 antibody using *Vangl2^-/-^* knock-out animals (*Vangl2^KO^*) (**Supp. Fig. 5C**) and a *Vangl2* knock-out cell line (Mentink et al., 2018). With this antibody, we observed that VANGL2 expression was enriched in epithelial cells in both mouse and human embryonic LV ChP (**Fig. 4E-G** and **Supp. Fig. 5D**). In contrast to a recent report (Langford et al., 2020), we found that VANGL2 exhibited a strong basolateral distribution typical for its role in mediating cell-to-cell signaling within the developing epithelium (Sittaramane et al., 2013). We also showed overlap of VANGL2 with the dedicated VANGL2 binding partner Scribble (Kallay et al., 2006), which was specifically restricted to the ChP epithelium during mouse and human embryonic development (**Supp. Fig. 5E, F**). Taken together, our data support a model in which VANGL2 is involved in the control of the WNT/PCP signaling in embryonic ChP.

**Figure 5.**
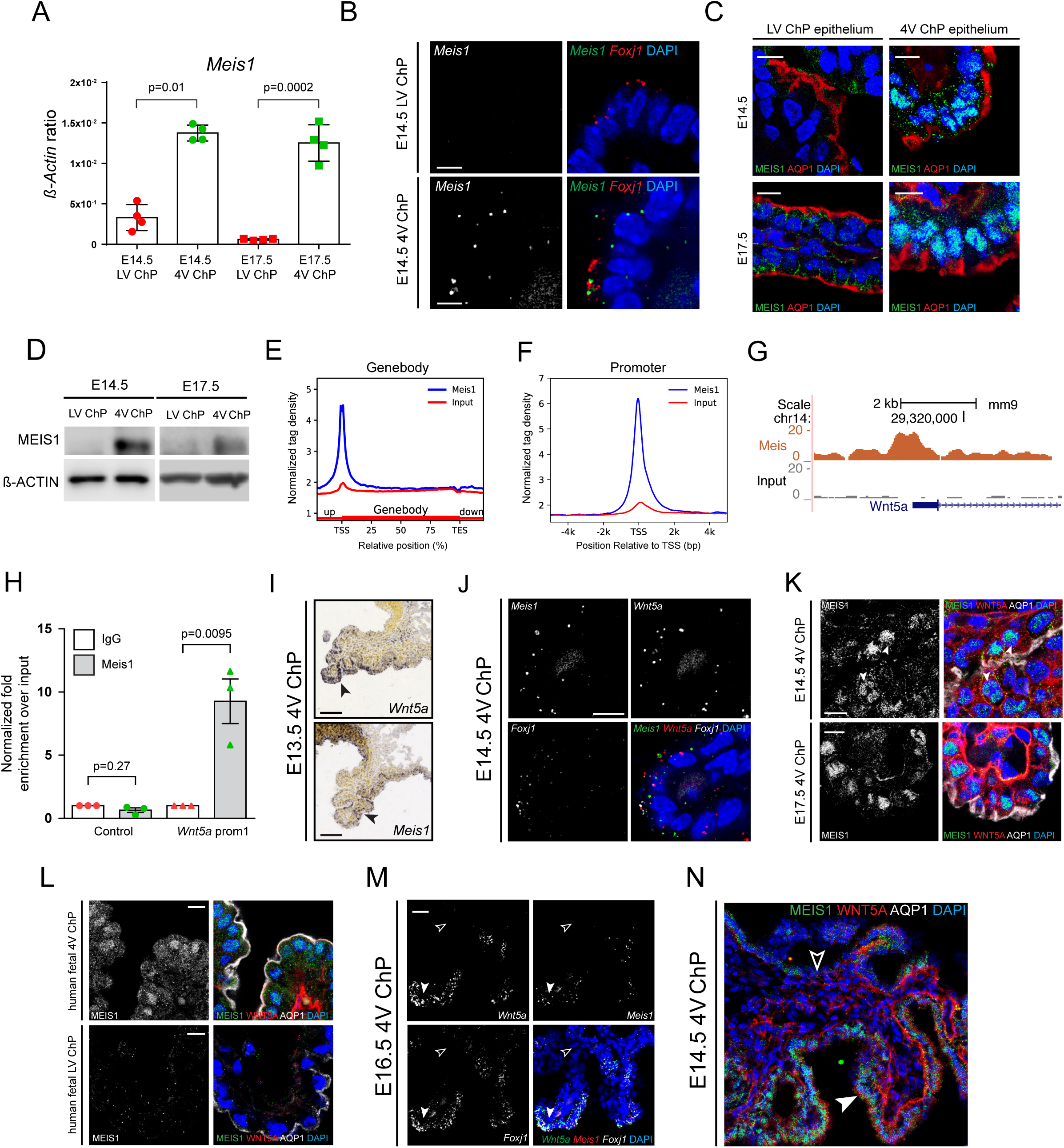
Meis1 expression pattern overlaps with Wnt5a expression in embryonic 4V ChP. (A) Real-time qPCR of *Meis1* expression in LV ChP and 4V ChP at E14.5 and E17.5. The expression was normalized against expression level of *β-actin* in each condition. Graph shows n=4 biologically independent samples; error bars represent mean ± s.d.; P values (two-tailed Student’s t-test with unequal variance). Biological replicates are indicated in the graph. (B) *In situ* hybridization analysis of the coronal sections showing *Meis1* expression being restricted to the 4V ChP epithelium and absent from the LV ChP epithelium at E14.5. *Foxj1* is a marker of ChP epithelium. Scale bar: 5 μ m. (C) Immunofluorescence analysis of MEIS1 in the LV ChP and 4V ChP at E14.5 and E17.5. Comparison with the AQP1, marker of ChP epithelium, shows absence of MEIS1 in the LV ChP, while MEIS1 can be detected in the 4V ChP epithelial layer. Scale bar: 10 μm. (D) Western blot analysis of MEIS1 protein in tissue lysates of LV ChP and 4V ChP. at E14.5 and E17.5. β-actin serves as a loading control. (E) MEIS1 density distribution over genebodies. The X-axis represents the relative position of the genebody. The Y-axis represents average MEIS1 and Input tag density over all Refseq genes. The 20% of up and downstream of genebody were extended based on corresponding genebody length. TSS, transcription start site, TES, transcription end site. (F) The X-axis represents the position relative to transcription start site (TSS), and Y- axis represents the average density of MEIS1 and Input. (G) A screenshot from the UCSC genome browser showing MEIS1 signal enriched in *Wnt5a* promoter. (H) The enrichment of Meis1 binding of Wnt5a promoter in E18.5 4V ChP with Cd19k36 enhancer used as control. Results are shown as relative fold enrichment over input in three independent experiments. (I) Sagittal sections showing similar *Meis1* and *Wnt5a* expression pattern in 4V ChP epithelium at E13.5. Image credit: Allen Institute. Scale bar: 100 μm. (J) *In situ* hybridization analysis of the coronal sections showing an overlapping pattern of expression of *Meis1* and *Wnt5a* within 4V ChP epithelium E14.5. *Foxj1* is a marker of ChP epithelium. Scale bar: 10 μm. (K) Immunofluorescence analysis of MEIS1 and WNT5A in 4V ChP epithelium at E14.5 and E17.5. AQP1 is a marker of ChP epithelium. Scale bar: 10 μm. (L) Immunofluorescence analysis showing MEIS1 detection being restricted only to 4V ChP epithelium as compared to LV ChP in 9-week-old human embryos. AQP1 is a marker of ChP epithelium. Scale bar: 10 μm. (M) *In situ* hybridization analysis of E16.5 coronal sections showing expression of *Meis1* and *Wnt5a* within the 4V ChP epithelium is restricted mostly to the tips of 4V ChP epithelium villi. *Foxj1* is a marker of ChP epithelium. Scale bar: 50 μm. (N) Immunofluorescence analysis showing the correlation of WNT5A signal intensity with MEIS1 signal within 4V ChP epithelium at E14.5 (arrowhead, empty arrowhead). Scale bar: 50 μm.

### Meis1 is expressed in *Wnt5a* expressing cells

We next sought to identify the mechanisms that control the expression of *Wnt5a* in the 4V ChP epithelium. The transcriptional co-activator Meis1 emerged as a likely candidate - Meis1 regulates *Wnt5a* expression in nearby branchial arches (Amin et al., 2015) and is highly expressed in the 4V ChP (Lun et al., 2015). We confirmed *Meis1* expression in 4V ChP throughout late embryogenesis (**Fig. 5A**) with *Meis1* transcripts being restricted to ChP epithelial cells marked by *Foxj1* (**Fig. 5B**).

We confirmed the presence of MEIS1 protein in 4V ChP from E14.5 to E17.5 (**Fig. 5C, D**) using validated antibodies against MEIS1 (**Supp. Fig. 6A, B**).

**Figure 6.**
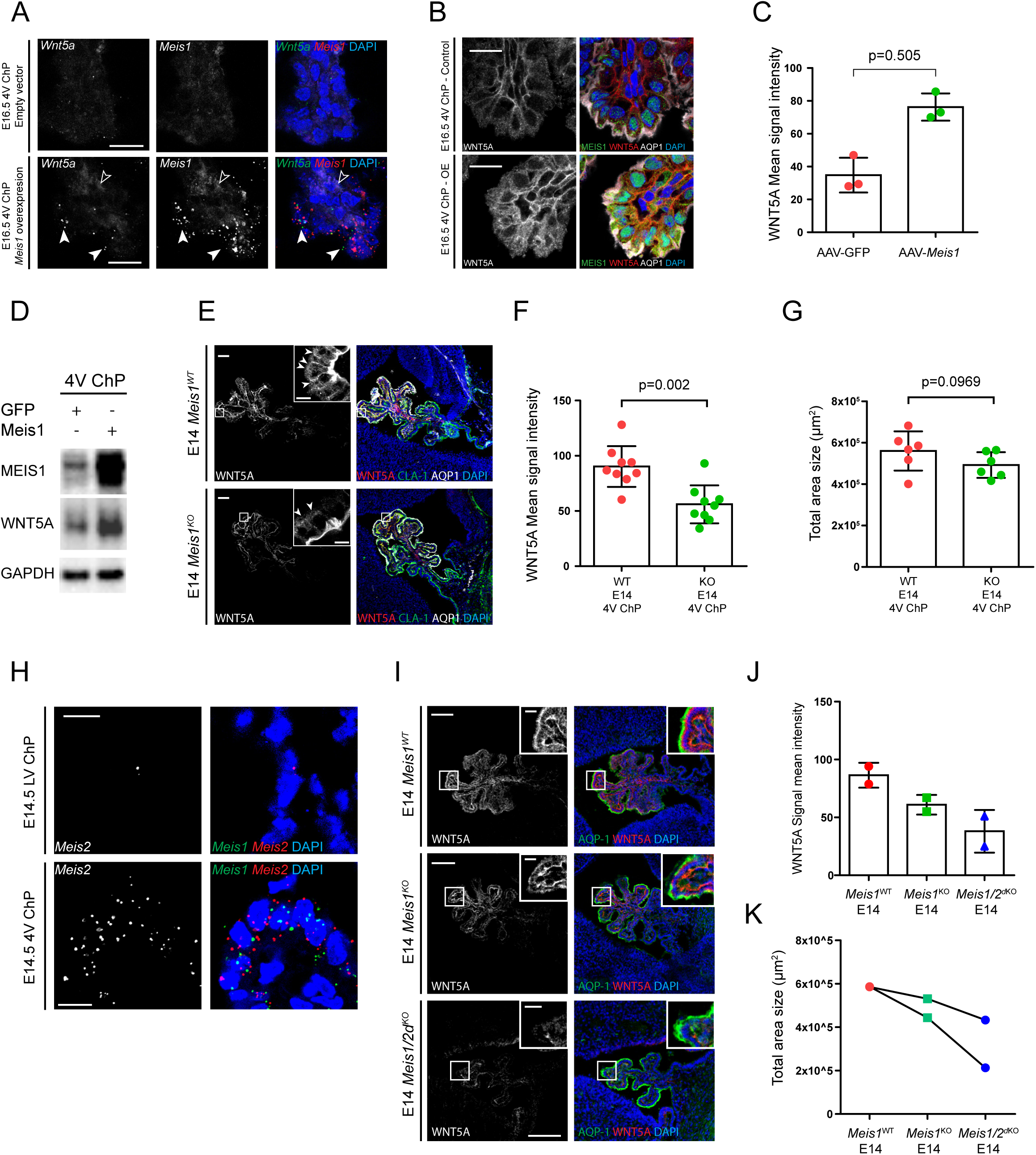
Meis1 regulates Wnt5a expression in 4V ChP epithelium. (A) *In situ* hybridization analysis of the coronal section showing *Wnt5a* expression in E16.5 4V ChP cells successfully transfected with *Meis1*-encoding vector (arrowheads) and low *Wnt5a* expression in untransfected cells (empty arrowheads). Scale bar: 20 μm. (B) Immunofluorescence analysis of MEIS1 and WNT5A signal in either control or *Meis1*-overexpressing E16.5 4V ChP epithelium targeted with a *Meis1*-encoding AAV vector shows upregulation of MEIS1 levels in 4V ChP epithelium that is associated with increased intensity of WNT5A as compared to 4V ChP epithelium transduced with control vector. AQP1 is a marker of ChP epithelium. Scale bar: 20 μm. (C) Quantification of mean WNT5A signal intensity difference between E16.5 AAV-GFP and AAV-*Meis1*-GFP transduced 4V ChP epithelium. Quantification has been performed using ImageJ software. Graph shows n=3 biologically independent samples; error bars represent mean ± s.d.; represented P values were obtained using a two-tailed Student’s t-test with unequal variance. Biological replicates are indicated in the graph. (D) Western blot analysis of MEIS1 and WNT5A protein levels in tissue lysates from AAV-GFP or AAV-*Meis1*-GFP transduced E16.5 4V ChP. Transduction with AAV-*Meis1*-GFP construct results in an increase of both MEIS1 and WNT5A protein levels in 4V ChP as compared to control vector. GAPDH serves as a loading control. (E) Immunofluorescence analysis of an embryonic sagittal section demonstrating a decrease of WNT5A signal in the E14 4V ChP epithelium of *Meis1^KO^* embryos as compared to *Meis1^WT^* embryonic 4V ChP (Insets; arrowheads). Inset images correspond to the highlighted boxed areas. Claudin-1 (Cla1) and AQP1 are markers of ChP epithelium. Scale bar: 50 μm. Inset scale bar: 10 μm. (F) Quantification of mean WNT5A signal intensity difference between E14 *Meis1^WT^* and *Meis1^KO^* 4V ChP epithelium. Quantification has been performed using ImageJ software. Graph shows n=9 biologically independent samples; error bars represent mean ± s.d.; represented P values were obtained using two-tailed Student’s t-test with unequal variance. Biological replicates are indicated in the graph. (G) Quantification of difference in total area size between E14 *Meis1^WT^* and *Meis1^KO^* 4V ChP epithelium. Quantification has been performed using ImageJ software. Graph shows n=6 biologically independent samples; error bars represent mean ± s.d.; represented P values were obtained using two-tailed Student’s t-test with unequal variance. Biological replicates are indicated in the graph. (H) *In situ* hybridization analysis of the coronal sections showing *Meis2* expression being restricted to 4th ChP epithelium and absent from LV ChP epithelium at E14.5 same as *Meis1*. *Foxj1* is a marker of ChP epithelium. Scale bar: 10 μm. (I) Immunofluorescence analysis of embryonic sagittal sections demonstrating difference in overall WNT5A levels (inset) and morphology between E14 *Meis1^WT^*, *Meis1^KO^* and *Meis1/2^dKO^* 4V ChP. AQP1 is a marker of ChP epithelium. Scale bar: 100 μm. Inset scale bar: 20 μm. (J) Quantification of mean WNT5A signal intensity difference between E14 *Meis1^WT^*, *Meis1^KO^* and *Meis1/2^dKO^* 4V ChP. Quantification has been performed using ImageJ software. Graph shows n=2 biologically independent samples; error bars represent mean ± s.d. (K) Quantification of difference in total area size between E14 *Meis1^WT^*, *Meis1^KO^* and *Meis1/2^dKO^* 4V ChP. Quantification has been performed using ImageJ software. Graph shows n=2 biologically independent samples; error bars represent mean ± s.d.

We identified the potential Meis1 binding site(s) upstream of *Wnt5a* by analyzing chromatin immunoprecipitation followed by sequencing (ChIP-seq) on E18.5 4V ChP. MEIS1 binding was enriched at the genomic region containing the proximal *Wnt5a* promoter (**Fig. 5E-G**). We confirmed Meis1 binding to the *Wnt5a* promoter by ChIP-qPCR (**Fig. 5H**). Consistent with these data, the spatial distribution of *Meis1* expression partially overlapped with *Wnt5a* expression in the 4V ChP (**Fig. 5I-J**, arrowheads). At the protein level, MEIS1 positive staining cells also displayed strong staining for WNT5A in both embryonic mouse and human 4V ChP epithelium (**Fig. 5K, L**), particularly at the distal tips of ChP villi (**Fig. 5M, N**). At later embryonic and early postnatal stages, the expression and protein levels of both MEIS1 and WNT5A progressively decreased in the 4V ChP, indicating a possible correlation between MEIS1 availability and expression level of *Wnt5a* (**Supp. Fig. 6C-E**).

### Meis1 controls expression of *WNT5A* in the embryonic 4V ChP

Next, we adopted an adeno-associated viral (AAV) approach using a serotype with tropism for ChP epithelial cells (Haddad et al., 2013) to investigate the effect of supplemental *Meis1* expression on 4V ChP development (**Supp. Fig. 7A, B**). We validated efficient intracerebroventricular delivery (at E13.5) and protein induction (by E16.5) by robust induction of EGFP signal in the epithelium of all the ChPs (**Supp. Fig. 7C, D**). We then successfully applied this approach to overexpress MEIS1 (**Supp. Fig. 7E-G**). *In situ* hybridization analysis showed *Wnt5a* upregulation that was restricted to the 4V ChP epithelial cells overexpressing *Meis1* (**Fig. 6A**) and that cells with the highest MEIS1 expression also exhibited the highest levels of WNT5A staining (**Fig. 6A, B** and **Supp. Fig. 7H**) with clear enrichment of overall WNT5A protein level (**Fig. 6C, D**).

**Figure 7.**
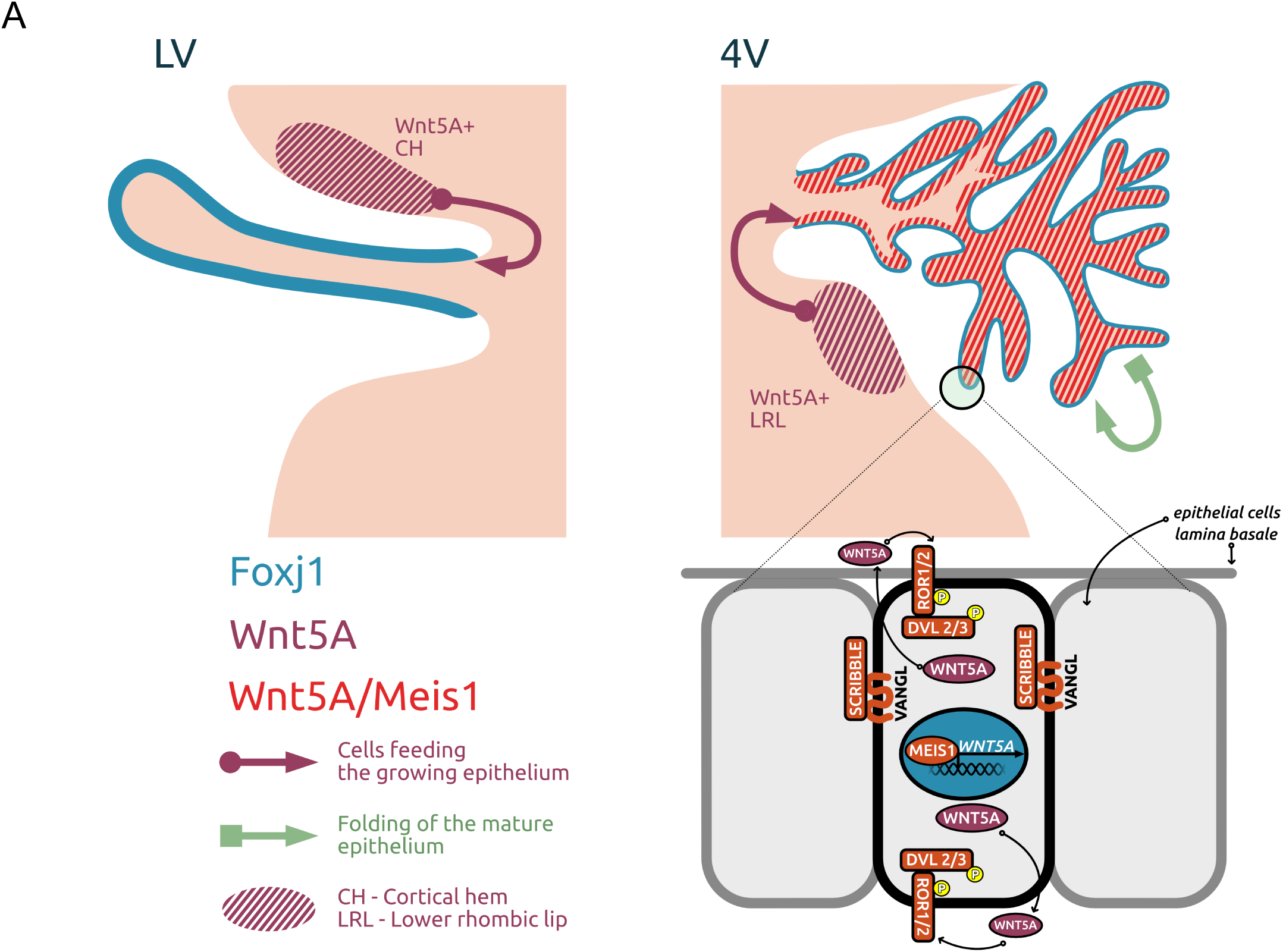
WNT5A signaling in the embryonic ChP epithelium. (A) Schematic depiction summarizing key aspects of WNT5A signaling in embryonic LV and 4V ChP epithelium. *Wnt5a* is expressed in region adjacent to both developing ChPs (CH for LV ChP and LRL for 4V ChP) with *Wnt5a* expression being restricted only to 4V ChP epithelium but not LV ChP epithelium. *Meis1* expression is also spatially restricted only to the 4V ChP epithelium as compared to LV ChP. *FoxJ1* is a marker gene for embryonic ChP epithelium. Epithelial WNT5A, whose production is under direct regulatory control of MEIS1 transcription co-factor, stimulates in an autocrine manner transmembrane receptors ROR1 and ROR2. Their activation leads to the activation of intracellular proteins DVL2 and DVL3. Ultimately, WNT5A signaling contributes to the suppression of WNT/canonical signaling, which is re-activated upon loss of WNT5A. Key components of the WNT/PCP signaling pathway, namely VANGL2 and its interacting partner SCRIBBLE also participate in the proper establishment of the ChP epithelium.

Based on the previously established role of *Wnt1* in 4V ChP development (Awatramani et al., 2003), we employed *Wnt1-Cre2* mouse line (Lewis et al., 2013) to conditionally delete *Meis1* (*Meis1^cKO^*) in the 4V ChP epithelium in order to functionally test the link between *Meis1* and *Wnt5a* expression (**Supp. Fig. 8A-B**).

We confirmed efficient reduction of *Meis1* expression in 4V ChP epithelial cells (**Supp. Fig. 6B**) but not in adjacent regions where *Wnt1* was not expressed (**Supp. Fig.8A, C**, arrowheads). Importantly, immunofluorescence analysis of *Meis1^cKO^* embryos revealed a prominent reduction of the WNT5A signal in the embryonic 4V ChP epithelium as compared to *Meis1^WT^* littermates (**Fig. 6E, F**). Unexpectedly, we did not observe any changes in overall 4V ChP morphology (**Fig. 6G**), which we hypothesized could be explained by having achieved only partial reduction of WNT5A levels and/or the possible redundant effects of other *Meis* factors including *Meis2*, which is also expressed in 4V ChP (Lun et al., 2015). To test this possibility, we first confirmed that *Meis2* expression was restricted to the developing 4V ChP epithelium similar to *Meis1* (**Fig. 6H**). Next, we conditionally deleted *Meis1* and *Meis2* (*Meis1/2^dKO^*) in the 4V ChP epithelium using *Meis1^cKO^* mouse line. *Meis1/2^dKO^* embryos were obtained with very low frequency. Comparing the ChP epithelium of two *Meis1/2^dKO^* embryos with control littermates showed *Meis2-*loss further decreased WNT5A protein levels in *Meis1/2^dKO^* embryos compared to *Meis1* knockout alone (**Fig. 6I, J**), supporting the model that *Meis1* and *Meis2* have additive effects on *Wnt5* expression. Notably, the two *Meis1/2^dKO^* embryos exhibited impaired morphogenesis and decreased overall size of the 4V ChP in comparison to WT and *Meis1^KO^* littermates (**Fig. 6K**). Taken together, these findings demonstrate a direct regulation of *Wnt5a* expression by MEIS1 in the developing 4V ChP and suggest redundancy of MEIS1 and MEIS2 in this process.

## DISCUSSION

Despite large overlap in their cellular architecture and secretory capacity, ChPs adopt dramatically different morphologies during development. Here we report a novel role for the epithelium-derived WNT5A in the regulation of tissue branching morphology within the ChP that is specific for the 4V. WNT5A mediated signaling, established as an important regulator of morphogenesis in various tissues (Yamaguchi et al., 1999), has been recently recognized to play an important role during ChP embryogenesis (Kaiser et al., 2019; Langford et al., 2020). Using temporally controlled conditional deletion of *WNT5A* directed to the ChP epithelium, we identified two roles of WNT5A, separated in time and space. Our data agree and expand upon recently published findings showing a crucial role of WNT5A in the development of embryonic ChPs (Langford et al., 2020).

By integrating information about WNT5A expression domains and functional data from various *WNT5A* mouse mutants, we propose a model in which WNT5A has two distinct roles in mouse ChP formation (**Figure 7**). Early on, *WNT5A* is expressed in the precursor domains that are directly adjacent to and feed the growth of 4V, 3V and LV ChP epithelium (Langford et al., 2020, Dani et al., 2019). Later on, *Wnt5a* is expressed in the maturing ChP epithelium in a spatially restricted fashion limited to the 4V ChP (Kaiser et al., 2019). Systemic *Wnt5a* deletion impairs development of all ChPs (Langford et al., 2020). In contrast, conditional deletion of WNT5A in time and space within FoxJ1-expressing epithelial cells (*Wnt5a^cKO^*) restricts the growth phenotype to the 4V ChP. The effect is age-dependent – strong when recombination is induced at E11.5 and non-detectable when induced at later stages. Our data show that the tamoxifen injection at E11.5 results in the loss of WNT5A protein at E12.5. Thus, WNT5A appears to be critical during this developmental period when branching of the 4V ChP begins. A similar temporal requirements for *Otx2* expression has also been described (Johansson et al., 2013).

We identify MEIS1 as one transcription factor that participates in the control of *Wnt5a* expression in the 4V ChP epithelium. Our data further suggest that MEIS1 and MEIS2 together regulate *Wnt5a* expression and 4V ChP development. Regionalized and conserved expression of *Wnt5a, Meis1,* and *Meis2* has previously been reported to occur during both human and mouse ChP embryogenesis (Kaiser et al., 2019, Lun et al., 2015). Nevertheless, despite our identification of this MEIS-WNT5A axis, the full transcriptional factor network that defines 4V ChP epithelium cell fate remains elusive. Potential co-regulation with HoxA2, which is also enriched in the 4V ChP epithelium (Awatramani et al., 2003; Lun et al., 2015), may be possible. HoxA2 binds the *Wnt5a* promoter (Donaldson et al., 2012), and in the branchial arches, Meis1 and HoxA2 synergize to positively regulate *Wnt5a* expression (Amin et al., 2015). Future studies may reveal if a similar evolutionary conserved mechanism contributes to the development of the ChP.

WNT5A controls cell polarity (Humphries and Mlodzik, 2018) as well as branching morphogenesis and tissue outgrowth in many developing tissues including mammary gland (Kessenbrock et al., 2017), kidney (Pietilä et al., 2016), lung (Li et al., 2005) and prostate gland (Huang et al., 2009). Similar effects have been described also in the developing midbrain (Andersson et al., 2008), and basolateral secretion of WNT5A by epithelial cells has been shown to participate in the process of lumen formation in kidney epithelium (Yamamoto et al., 2015). These findings point to a more universal role of epithelial WNT5A in the regulation of branching and outgrowth of epithelial sheets. Consistent with this model, we detected regionally restricted activation of non-canonical Wnt/PCP signaling in the ChP, with higher activity in the 4V ChP compared to the LV ChP. Importantly, WNT5A appears to be necessary for the initial stages of ChP development, but it is not essential after E14 despite continuing 4V ChP tissue growth and folding that progresses with development.

The function of WNT5A in the 4V ChP is analogous to its function in the intestinal epithelium, which also develops in a conveyor belt-like mechanism. Recent findings show a link between altered levels of WNT5A and perturbed development of the small intestine, largely resembling effects observed in the developing 4V ChP upon WNT5A deletion. Notably, either WNT5A deficiency or overexpression leads to similar phenotypes in both tissues (van Amerongen et al., 2012, Cervantes et al., 2009). Tightly controlled WNT5A levels thus seem to be essential for proper epithelial folding of both the 4V ChP and intestine and may represent a universal mechanism of epithelial development.

WNT5A is known to contribute to the balance between Wnt/β-catenin (canonical) and non-canonical Wnt signaling that is essential for proper tissue morphogenesis and homeostasis (Alexander et al., 2012). Crosstalk between WNT5A and other Wnts may differ for the two expression domains of *Wnt5a* in the developing ChP (see **Figure 7**). As *Wnt5a* expression domains are located adjacent to all embryonic ChPs (Langford et al., 2020) and overlap with the expression domains of Wnt ligands that activate β-catenin pathway, it is possible that *Wnt5a* acts in concert with other Wnt genes. For example, with the typical canonical ligand WNT2b, that is known to be an important patterning factor orchestrating proper development of ChPs in the cortical hem (Grove et al., 1998). Importantly, the Wnt5a domain also overlaps with an R-spondin (R-SPO)-positive progenitor domain (Dani et al., 2019). R-SPO acts as the amplifier of the Wnt/ β-catenin signaling which suggests that in the progenitor domain WNT5A can synergize or coordinate with Wnt/β-catenin signals. In contrast, in the 4V in the ChP epithelium WNT5A seems to be a signaling factor suppressing canonical WNT signaling as reported in mammary gland development (Roarty et al., 2009). Higher WNT/β-catenin signaling (as shown by upregulation of target genes *Axin2* or *Lef1* or phosphorylation of LRP6) was detected in the embryonic LV ChP epithelium with low/undetectable *Wnt5a* expression or upon deletion of *Wnt5a* in the 4V ChP. Notably, in comparison to *Wnt5a,* both *Axin2* or *Lef1* show a reverse pattern of expression along the P-D axis in the embryonic 4V ChP. This observation suggests that the spatial control of WNT/β-catenin signaling is achieved by increasing levels of WNT5A within the ChP epithelium (Sato et al., 2010).

In summary, our data reveal the complex functions of WNT5A in regulating ChP development. From a clinical perspective, rare neoplasms of the ChP have higher incidence in children and display regional specificity (Sun et al., 2014). Thus, elucidating the regionalized signaling pathways underlying ChP development may expose molecular mechanisms that lend one ventricle more or less susceptible to cancer. Because the ChP represents a tissue amenable to genetic manipulation (Chen et al., 2020), and it adopts distinct, ventricle-specific morphologies due to inherent differences in activation of WNT5A-driven PCP signaling, the ChP provides a tractable model for future research investigating the mechanistic basis of WNT5A signaling in tissue morphogenesis.

## SUPPLEMENTARY FIGURES

**Supplementary Figure 1.**
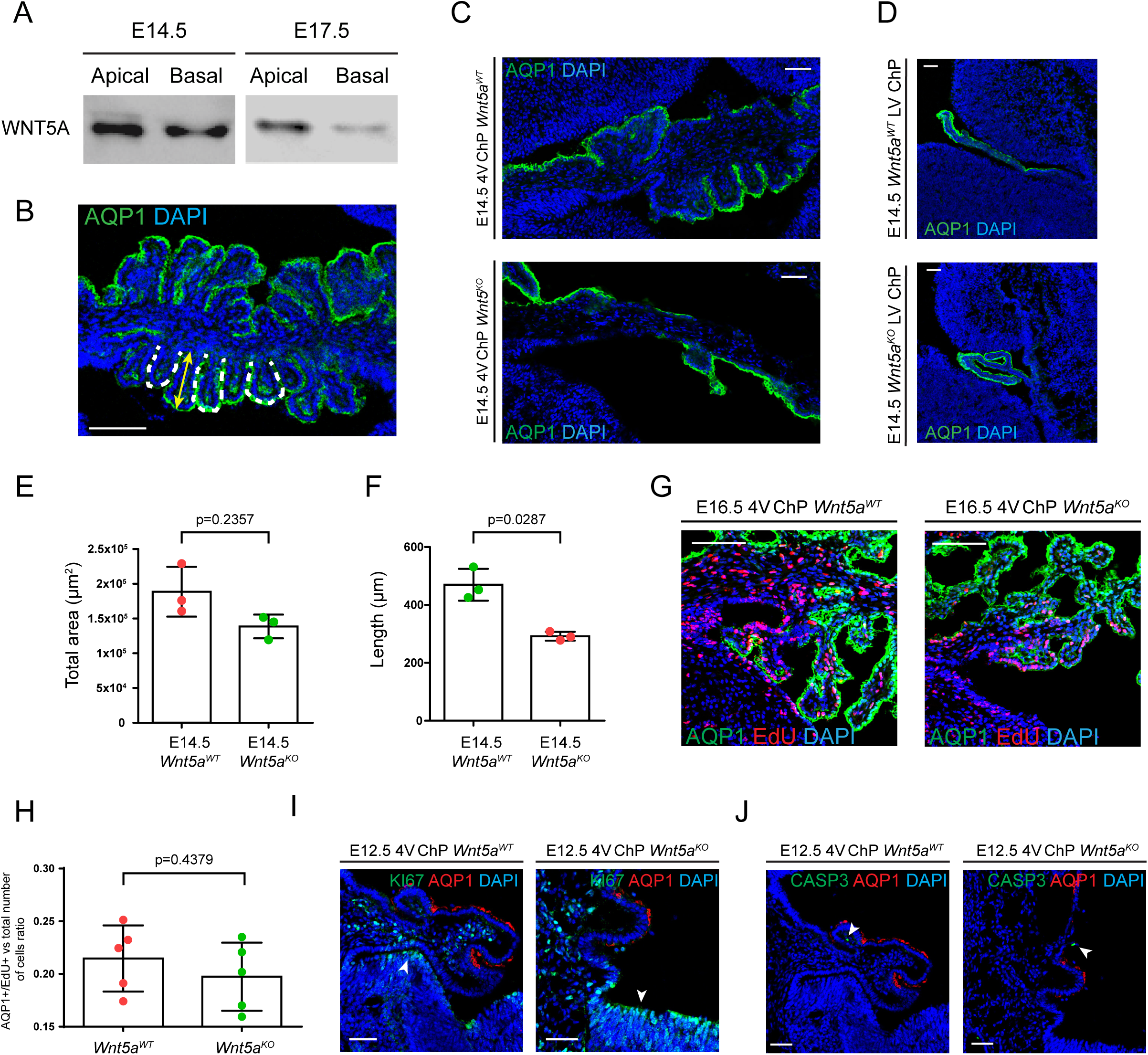
Wnt5a ablation during 4V ChP embryogenesis does not affect proliferation or apoptosis of the ChP epithelium. (A) Western blot analysis of transwell ChP-epithelial primary cultures derived from LV ChP and 4V ChP isolated from E14.5 and E17.5 embryos show bi-directional secretion of WNT5A to both apical and basal compartments. (B) Representative image of E16.5 4V ChP coronal section illustrating the process of branching point quantification and measurement of their length as used for the statistical analysis of the effect of *Wnt5a* ablation on 4V ChP embryogenesis. Dotted line highlights a representative villus as used for the analysis of branching morphology. Yellow double arrowhead line indicates total length of epithelial villi used for the quantification analysis. AQP1 is a marker of ChP epithelium Scale bar: 100 µm. (C) Representative image of E14.5 4V ChP coronal section showing altered overall morphology of the tissue between *Wnt5a^WT^* and *Wnt5a^KO^* embryos. AQP1 is a marker of ChP epithelium Scale bar: 100 µm. (D) Immunofluorescence analysis of E14.5 coronal section of LV ChP in *Wnt5a^WT^* and *Wnt5a^KO^* embryos demonstrates the aberrant morphology of the tissue. AQP1 is marker of ChP epithelium. Scale bar: 50 µm. (E, F) Statistical assessment of E16.5 LV ChP morphology shows there are no changes in the total area size (I) but a significant difference in the average branch length (J) between *Wnt5a^WT^* vs *Wnt5a^KO^* embryos. Graph shows n=3 biologically independent samples; error bars represent mean ± s.d.; P values (two-tailed Student’s t-test with unequal variance). Biological replicates are indicated in the graph. (G) Representative image of EdU staining in *Wnt5a^WT^* and *Wnt5a^KO^* embryonic 4V ChP at E16.5 shows no significant difference in the number of EdU+ cells incorporated within the epithelial layer. EdU injection was administered at E13.5 and embryos were collected 72h later. AQP1 is a marker of ChP epithelium Scale bar: 50 µm. (H) Statistical confirmation of lack of difference in the total number of EdU+ cells within the epithelium. Only EdU+/AQP1+ cells were used for analysis. Quantification has been performed by ImageJ software. Graph shows n=5 biologically independent samples; error bars represent mean ± s.d.; P values (two-tailed Student’s t-test with unequal variance). Biological replicates are indicated in the graph. (I) Immunofluorescence analysis of proliferation shows no difference in the proliferation marker KI67 signal distribution in 4V ChP epithelium or nearby progenitor zone (arrowheads) between *Wnt5a^WT^* and *Wnt5a^KO^* mouse embryos at E12.5. AQP1 is a marker of ChP epithelium. Scale bar: 50 µm. (J) Immunofluorescence analysis of apoptotic marker Caspase-3 (CASP3) demonstrates no difference in apoptotic activity with very few CASP3+ cells (arrowheads) being seen in the 4V ChP epithelium in both *Wnt5a^WT^* and *Wnt5a^cKO^* mouse embryos at E14.5. AQP1 is a marker of ChP epithelium. Scale bar represents 50 µm.

**Supplementary Figure 2.**
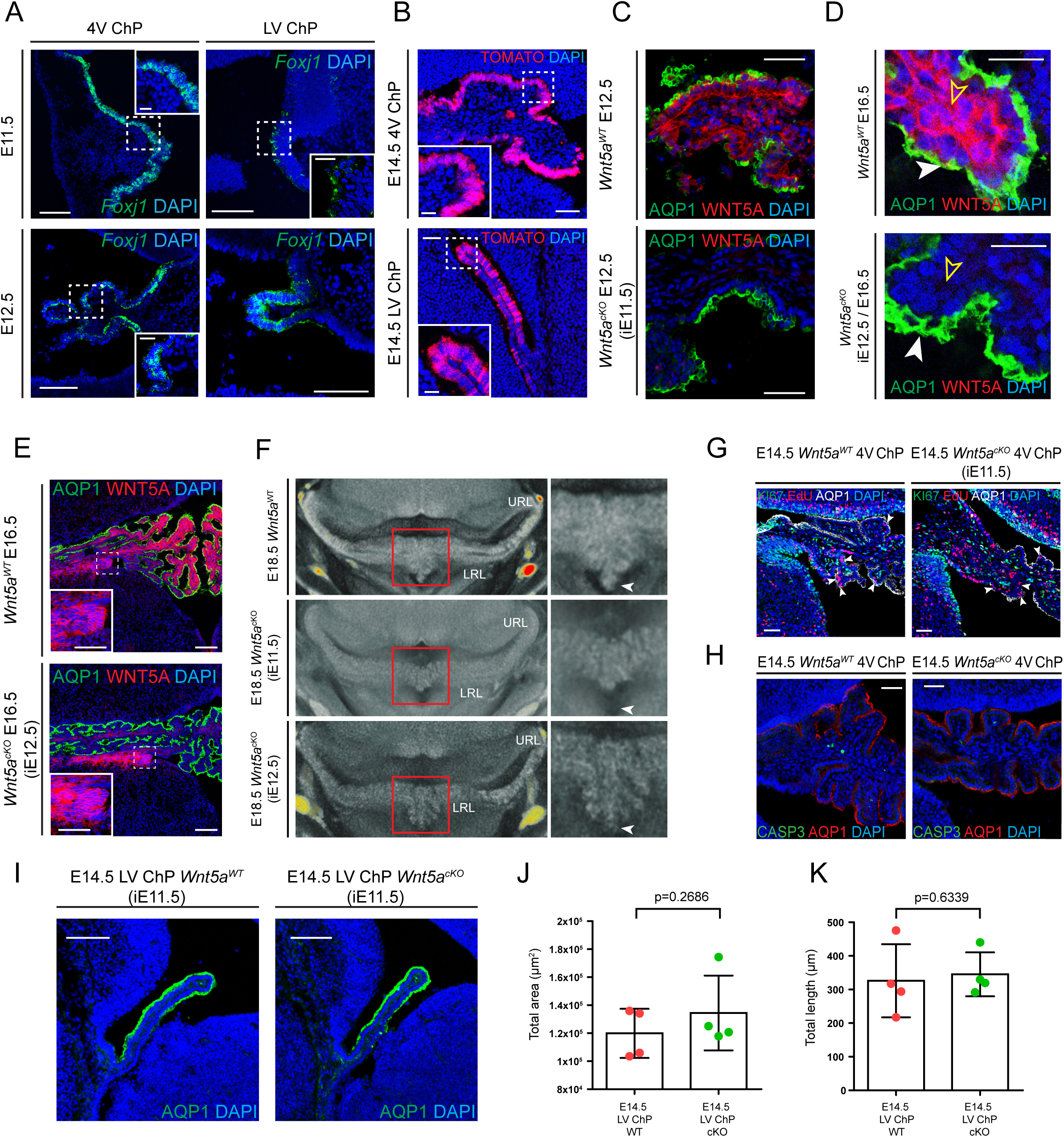
Morphological effects of Wnt5a ablation on 4V ChP development persist to later embryogenic stages. (A) *In situ* hybridization analysis of the coronal section showing expression pattern of *Foxj1* in 4V ChP epithelium at E11.5 and E12.5 which were the time points used for the generation of the ChP-specific ablation of *Wnt5a* in *Wnt5a^cKO^* embryos. Inset highlights detection of *Foxj1* expression restricted to ChP epithelium. Scale bar: 100 μm. Inset scale bar: 20 μm. (B) Immunostaining images of coronal sections from E14.5 *FoxJ1-CreERT2-tdTomato* embryos with tamoxifen injection at E10.5. Inset image highlights specificity of recombination restricted to 4V ChP and LV ChP epithelium as tracked by tdTomato staining. Scale bar: 50 µm. Inset scale bar: 20 µm. (C) Immunofluorescence analysis of WNT5A in 4V ChP in *Wnt5a^cKO^* (with tamoxifen injection at E11.5) and *Wnt5a^WT^* embryos at E12.5. WNT5A signal detection in completely absent from *Wnt5a^cKO^* 4V ChP epithelium as compared to *Wnt5a^WT^* embryo. AQP1 is a marker of ChP epithelium. Scale bar: 50 µm. (D) Immunofluorescence analysis of E16.5 4V ChP epithelium confirms the epithelial origin (arrowhead) of the WNT5A detected in the stromal compartment of the tissue (yellow empty arrowhead) as the WNT5A staining is completely abolished in *Wnt5a^cKO^* embryonic 4V ChP epithelium. AQP1 is marker of ChP epithelium Scale bar: 10µm. (E) Immunofluorescence analysis of WNT5A in 4V ChP in *Wnt5a^cKO^* and *Wnt5a^WT^* embryos at E16.5. Inset (dotted line box) detailed view of WNT5A signal detected in both *Wnt5a^cKO^* and in *Wnt5a^WT^* in the domain directly adjacent to 4V ChP. Inset (dotted line box) showing that WNT5A signal detection is completely absent from *Wnt5a^cKO^* 4V ChP epithelium as compared to *Wnt5a^WT^* embryo. AQP1 is a marker of ChP epithelium. Scale bar: 100 µm. Inset scale bar: 50 µm. (F) Transverse sections of mouse embryonic brain at E18.5 using the microCT method shows the hypomorphic phenotype, most pronounced in the ventral region of 4th ChP, in *Wnt5a^cKO^* embryos with tamoxifen induction at E11.5 as compared to *Wnt5a^WT^* or *Wnt5a^cKO^* embryos (arrowheads). In contrast, *Wnt5a^cKO^* embryos with tamoxifen induction at E12.5 display a 4th ChP morphological phenotype that is comparable to the 4V ChP *Wnt5a^WT^* littermates. (G) Representative image of EdU staining in *Wnt5a^WT^* and *Wnt5a^cKO^* (tamoxifen induction at E11.5) embryonic 4V ChP at E14.5 shows no significant difference in the amount of EdU+ cells incorporated within epithelial layer (arrowheads). EdU injection was administered at E11.5 and embryos were collected 72h later. AQP1 is marker of ChP epithelium. Scale bar: 50 µm. (H) Immunofluorescence analysis of CASP3 demonstrates no difference in the number of apoptotic cells in 4V ChP epithelium between *Wnt5a^WT^* and E11.5 induced *Wnt5a^cKO^* mouse embryos at E14.5. Scale bar: 50 µm. (I) Immunofluorescence analysis E14.5 LV ChP coronal section shows absence of morphological difference between *Wnt5a^WT^* and E11.5 mouse embryos. AQP1 is marker of ChP epithelium. Scale bar: 50 µm. (J, K) Statistical assessment of E14.5 LV ChP morphology shows no difference in the LV ChP in terms of the total area size (J) and average length (K) between *Wnt5a^WT^* vs *Wnt5a^cKO^* embryos (recombination induced at E11.5). Graph shows n=4 biologically independent samples; error bars represent mean ± s.d.; P values (two-tailed Student’s t-test with unequal variance). Biological replicates are indicated in the graph.

**Supplementary Figure 3.**
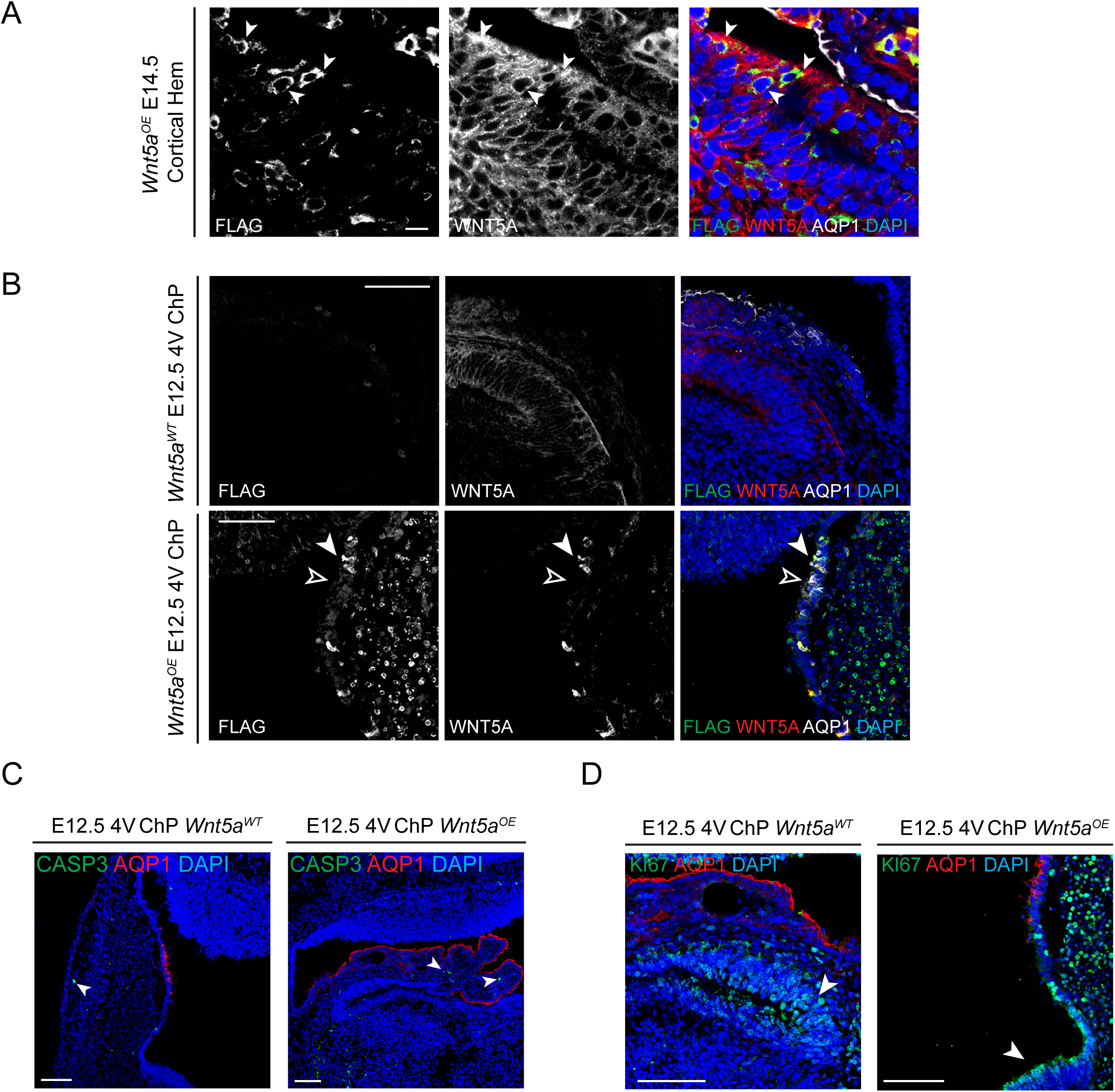
Embryonic 4V ChP morphological phenotype observed in *Wnt5a^OE^* embryos is not associated with overt changes in proliferation or apoptosis. (A) Immunofluorescence analysis of E14.5 coronal sections showing the presence of ectopically expressed WNT5A-FLAG in the cortical hem region (arrowheads) located adjacent to the developing LV ChP. AQP1 is marker of ChP epithelium. Scale bar: 10 µm. (B) Immunofluorescence analysis of E12.5 coronal sections showing induction of FLAG-WNT5A (arrowhead) overexpression in cells adjacent to AQP1+ epithelial cells (empty arrowhead) of 4V ChP in *Wnt5a^OE^* embryos as compared to Wnt5aWT littermates. AQP1 is a marker of ChP epithelium. Scale bar: 100 μm. (C) Immunofluorescence analysis of E12.5 coronal sections demonstrates no difference in Caspase-3 (CASP3) signal distribution indicating comparably low levels of apoptotic cell death in 4V ChP, which is mostly restricted to the stromal compartment of the tissue (arrowheads), between *Wnt5a^WT^* and *Wnt5a^OE^* mouse embryos. AQP1 is a marker of ChP epithelium. Scale bar: 100 µm. (D) Immunofluorescence analysis of E12.5 coronal sections does not reveal any significant change in KI67 signal distribution indicating comparable proliferative activity in the progenitor domain adjacent to 4V ChP between *Wnt5a^WT^* and *Wnt5a^OE^* mouse embryos. AQP1 is marker of ChP epithelium. Scale bar: 100 µm.

**Supplementary Figure 4.**
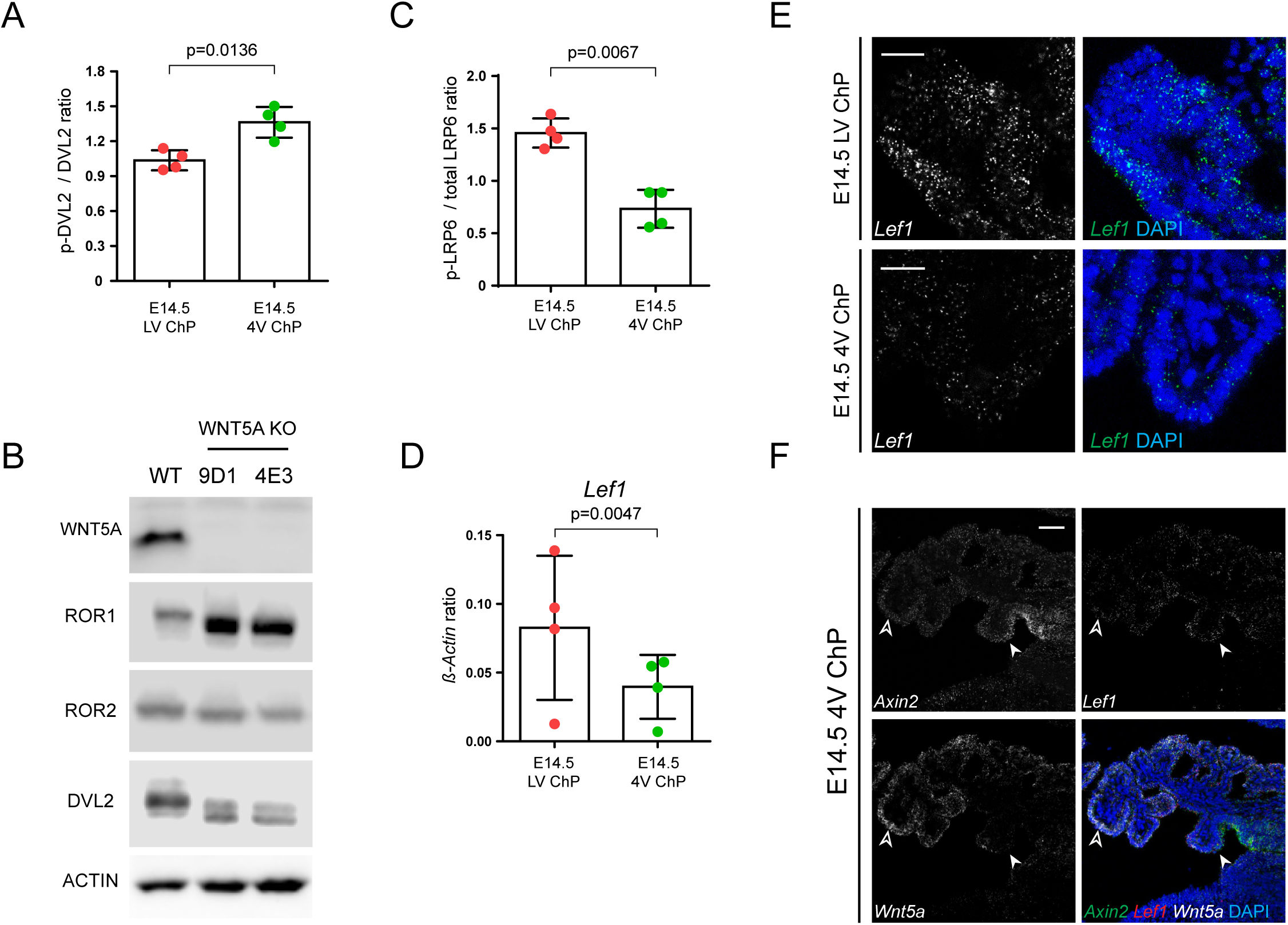
WNT5A regulates the activation of canonical and non-canonical signaling. (A) Statistical analysis of immunoblot p-DVL2 / DVL2 protein levels. Graph shows n=4 biologically independent samples; error bars represent mean ± s.d.; P values (two-tailed Student’s t-test with unequal variance). Biological replicates are indicated in the graph. (B) Western blot analysis of *Wnt5a^WT^* and *Wnt5a^KO^* clones derived from HEK293T cell line shows loss of ROR1, ROR2, DVL2 and DVL3 shift upon WNT5A ablation validating their role as a specific readout of WNT5A-mediated signaling activity. β-actin serves as a loading control. (C) Statistical analysis of immunoblot p-LRP6 / total LRP6 protein levels. Graph shows n=4 biologically independent samples; error bars represent mean ± s.d.; P values (two-tailed Student’s t-test with unequal variance). Biological replicates are indicated in the graph. (D) Real-time qPCR of Wnt canonical target gene *Lef1* expression in LV ChP and 4V ChP at E14.5. The expression was normalized against the expression level of *β-actin* in each condition. Graph shows n=4 biologically independent samples; error bars represent mean ± s.d.; P values (two-tailed Student’s t-test with unequal variance). Biological replicates are indicated in the graph. (E) *In situ* hybridization analysis of coronal sections confirming increased *Lef1* expression in E14.5 LV ChP epithelium as compared to 4V ChP epithelium. Scale bar: 20 μm. (F) *In situ* hybridization analysis of coronal sections showing gradual decrease of *Axin2* and *Lef1* (arrowhead) expression levels along the lateral-medial axis of the 4V ChP epithelium. This decrease is associated with an increase in the expression of *Wnt5a* (empty arrowhead) in the medial direction within the 4V ChP epithelium. Scale bar: 50 μm.

**Supplementary Figure 5.**
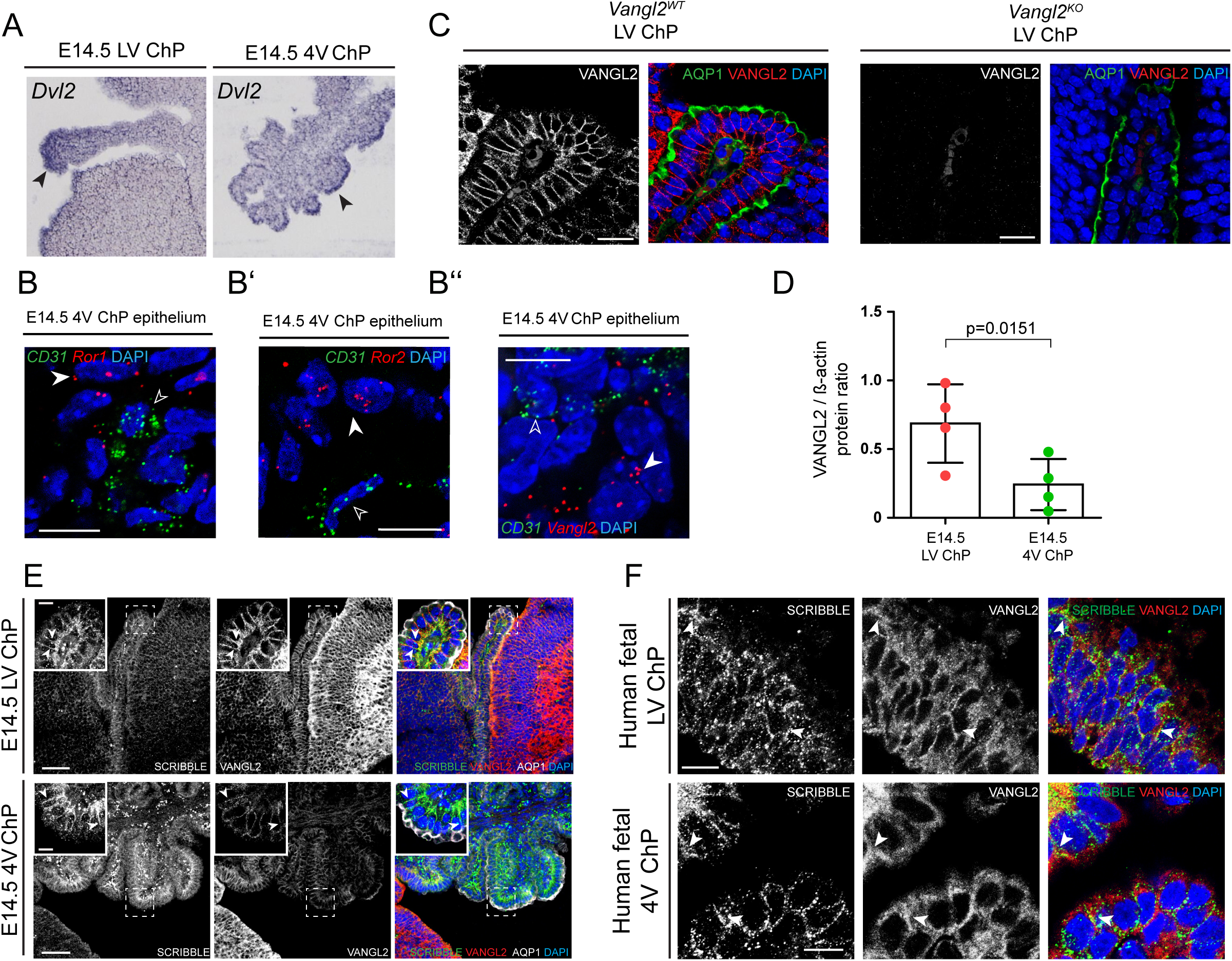
Non-canonical Wnt signaling components are located in the embryonic ChP epithelium. (A) Sagittal sections showing the *Dvl2* expression pattern in LV and 4V ChP epithelium at E14.5 with increased expression levels detected in ChP epithelium (arrowhead). Data are adopted from http://www.eurexpress.org/ee/. (B, B’, B’’) *In situ* hybridization analysis of the coronal sections of E14.5 4V ChP demonstrates an absence of the expression of non-canonical Wnt pathway downstream components (arrowheads) *Ror1* (B), *Ror2* (B’) and *Vangl2* (B’’) from the stromal compartment as highlighted by the expression of CD31 (empty arrowhead), marker of endothelial cells. Scale bar: 10 μm. (C) Immunofluorescence analysis of E14.5 coronal section showing the loss of VANGL2 staining in the LV ChP epithelium in *Vangl2^KO^* as compared to *Vangl2^WT^* embryos. AQP1 is a marker of ChP epithelium. Scale bar: 20 μm. (D) Statistical analysis of immunoblot VANGL2 protein level. Graph shows n=4 biologically independent samples; error bars represent mean ± s.d.; P values (two-tailed Student’s t-test with unequal variance). Biological replicates are indicated in the graph. (E) Immunofluorescence analysis of E14.5 LV and 4V ChP coronal sections showing signal detection overlap for VANGL2 and the scribble homolog (SCRIBBLE) protein that is restricted mostly to the ChP epithelial layer (Inset). Arrowheads highlight overlap of SCRIBBLE and VANGL2 within the basolateral domain of the epithelium. AQP1 is a marker of ChP epithelium. Scale bar: 50 μm. Inset scale bar: 10 μm. (F) Immunofluorescence analysis of a 9-week-old human fetal LV and 4V ChP coronal sections showing signal detection overlap for VANGL2 and hScrib (SCRIBBLE) protein in the ChP epithelial layer. Arrowheads highlight overlap of SCRIBBLE and VANGL2 within the basolateral domain of the epithelium. AQP1 is a marker of ChP epithelium. Scale bar: 10 μm.

**Supplementary Figure 6.**
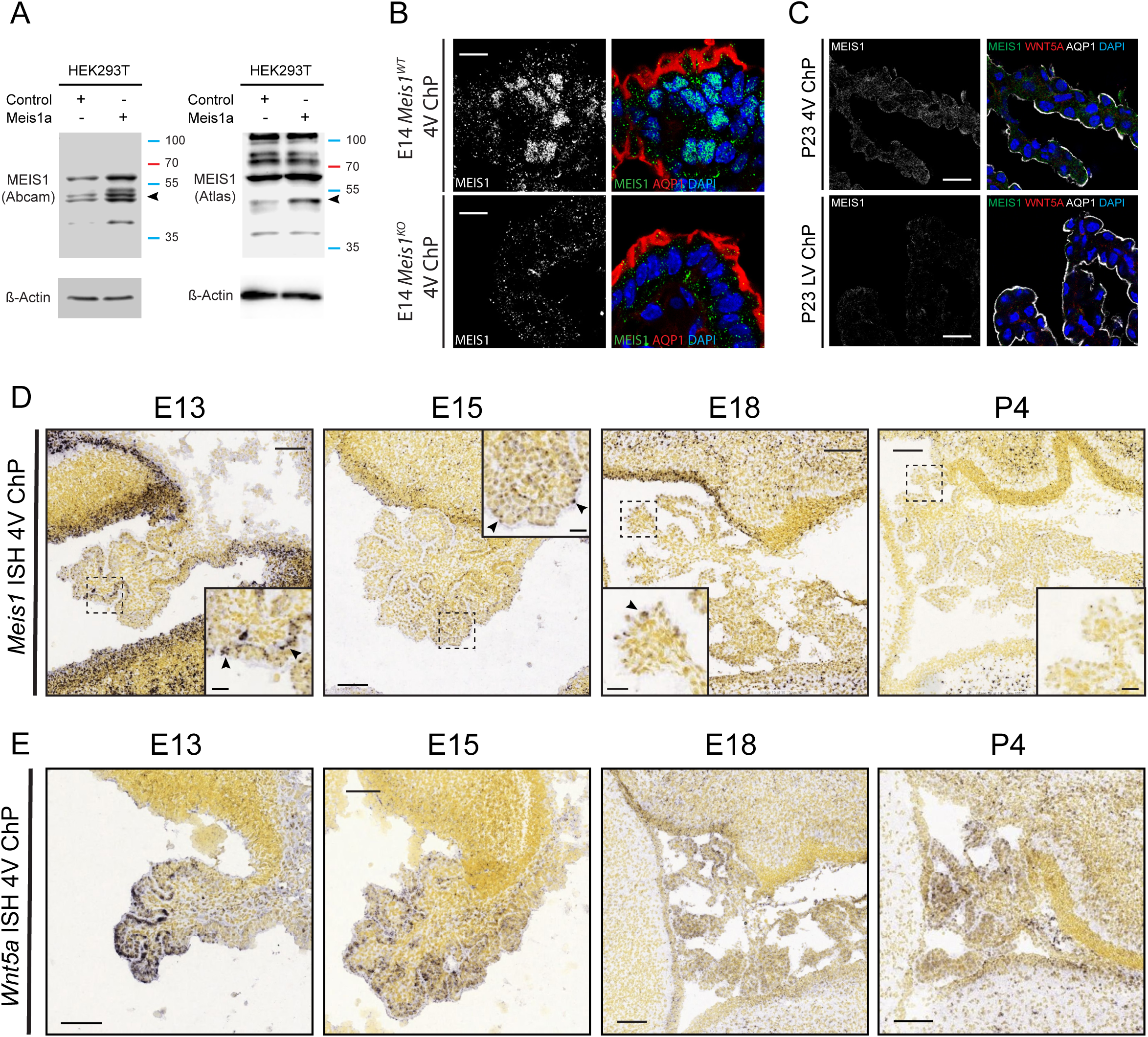
Meis1 expression in 4V ChP is reduced at postnatal stages. (A) Western blot validation of MEIS1 antibodies (Abcam - cat.no. 19867; Atlas - ncat.no HPA056000) in cell lysates of HEK293T cells overexpressing *Meis1a* encoding construct or empty control plasmid. β-actin serves as a loading control. Arrowhead indicating specific band corresponding to the expected protein size of MEIS1 (43 kDa - source: uniprot.org). (B) Immunofluorescence analysis of embryonic coronal sections showing the loss of signal for MEIS1 in the 4V ChP epithelium of *Meis1^KO^* embryos as compared to *Meis1^WT^* animals embryonic 4V ChP at E14.5. AQP1 is marker of ChP epithelium. Scale bar: 10 μm. (C) Immunofluorescence analysis of P23 coronal sections showing loss of MEIS1 and WNT5A signal detection in both LV and 4V ChP epithelium. AQP1 is a marker of ChP epithelium. Scale bar: 20 μm. (D) Sagittal sections showing a similar gradual decrease of *Meis1* expression during postnatal stages in 4V ChP. Magnified view highlights the expression of *Meis1* restricted to the 4V ChP epithelium (arrowheads). Image credit: Allen Institute. Scale bar: 100 μm. Inset scale bar: 25 μm. (E) Sagittal sections showing similar gradual decreases of *Wnt5a* expression during postnatal stages in 4V ChP. Image credit: Allen Institute. Scale bar: 100 μm.

**Supplementary Figure 7.**
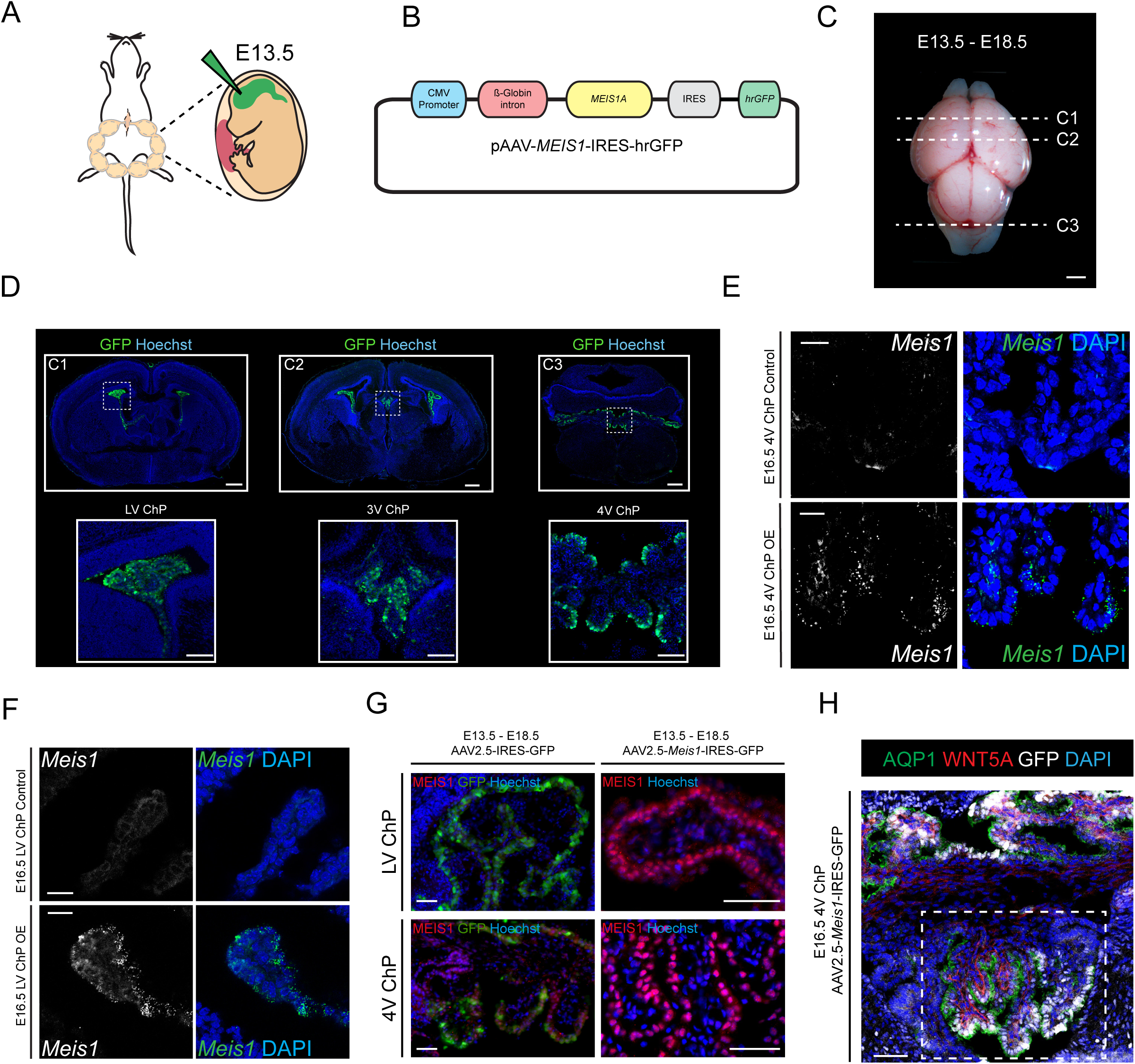
Validation of AAV-mediated Meis1 overexpression in embryonic ChP. (A) Schematic of intracerebroventricular associated adeno-associated viral (AAV) particle delivery. (B) Schematic of AAV-*Meis1* construct. (C) Schematic of anatomical localization of embryonic ChPs at E18.5. Scale bar: 1 mm. (D) Immunofluorescent analysis of coronal sections showing efficient targeting of embryonic ChP epithelium using AAV-based delivery system. Inset showing higher magnification of embryonic choroid plexuses, C1-LV ChP, C2 - 3V ChP and C3 - 4V ChP as outlined in (C). Scale bar: 100 μm. Inset scale bar: 30 μm (E) *In situ* hybridization analysis of E16.5 4V ChP coronal section showing efficient induction of *Meis1* expression in epithelial cells upon transfection with AAV-*Meis1* construct. Scale bar: 20 μm. (F) *In situ* hybridization analysis of E16.5 LV ChP coronal section showing efficient induction of *Meis1* expression in epithelial cells upon transfection with AAV-*Meis1* construct. Scale bar: 20 μm. (G) Immunofluorescence analysis of E18.5 coronal sections showing of the efficient induction of MEIS1 protein production in both LV and 4V ChP epithelium upon transduction with AAV-*Meis1* encoding construct, while no MEIS1 signal can be detected in LV ChP transduced with empty AAV vector. Scale bar: 100 μm. (H) Immunofluorescence analysis of E16.5 4V ChP coronal section illustrating the area (dotted box) used for WNT5A signal intensity quantification. Scale bar: 50 µm.

**Supplementary Figure 8.**
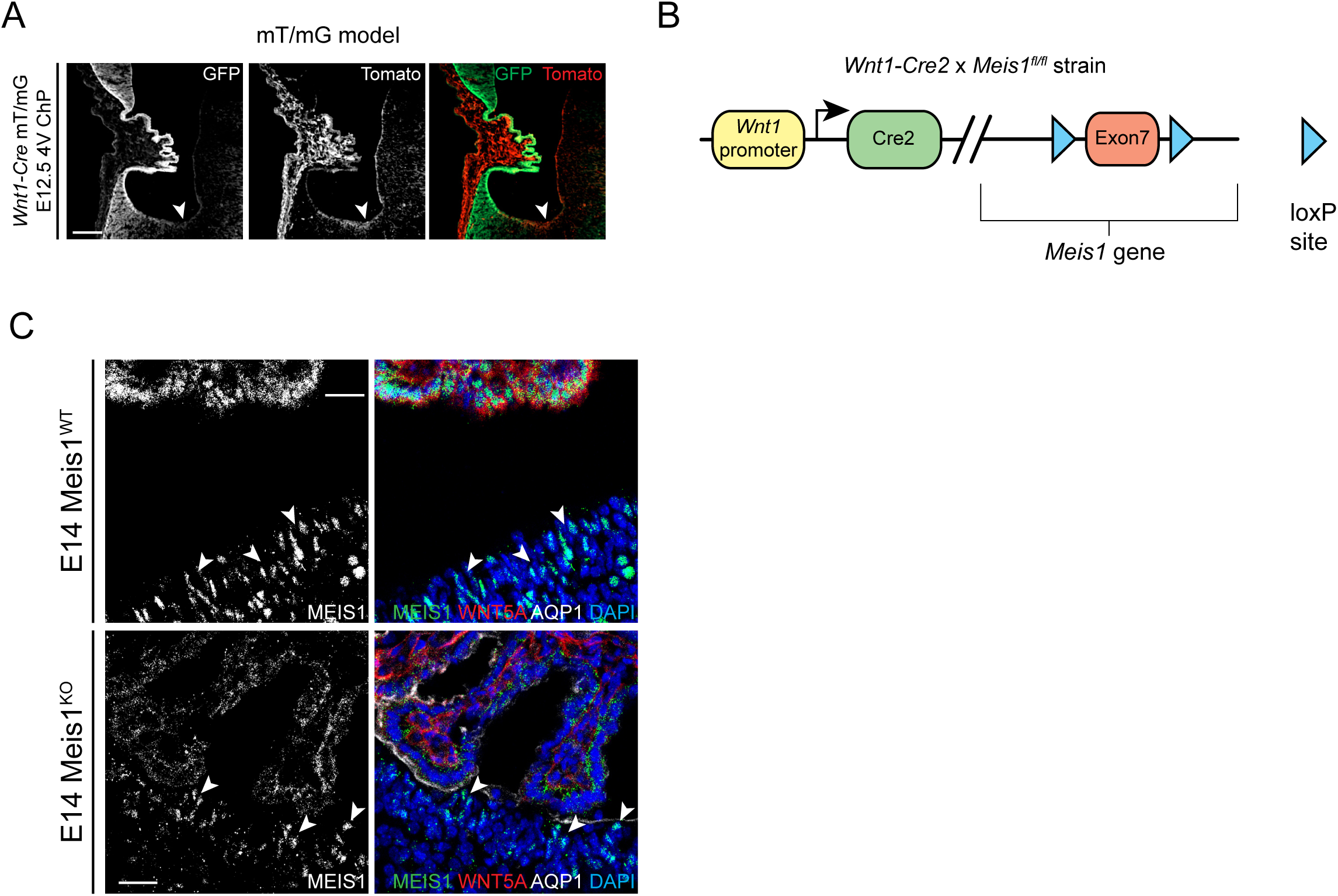
Meis1 conditional knock-out mouse strain validation. (A) Immunofluorescence analysis of an E12.5 4V ChP sagittal section showing a specific pattern of Wnt1 promoter-driven recombination activity that is restricted to the ChP epithelium and absent from stromal compartment. Arrowhead highlights region displaying absence of Wnt1-driven recombination Scale bar: 100 μm. (B) Schematic of Wnt1-Cre x *Meis1^flox/flox^* (*Meis1^cKO^*) mouse strain generation. (C) Confirmation of spatial selectivity of *Meis1* ablation specific for the 4V ChP epithelium as compared to adjacent floor plate (arrowheads). AQP1 is a marker of ChP epithelium. Scale bar: 25 μm.

## METHODS

### Mouse strains

Conditional *Meis1*^fl/fl^ mice were generated from the embryonic stem cell clone HEPD0632_4_H07 purchased from EUCOMM. The Frt-flanked LacZ/neo cassette was removed by ACTFLPe (strain #005703). LoxP sites flank exon ENSMUSE00000655363 encoding the homeobox region of the *Meis1* gene. Genotyping primers: CTGCGCTTCCTACATCACTG and CACTTCAGCGTCACTTGGAA produce 227-bp fragment for the wild-type allele and 262-bp band for the floxed allele.

Conditional *Meis2 ^fl/fl^* strain with loxP sites around exons 2-6 was described earlier (Machon et al., 2015). Genotyping primers: GCAAGGGTGCTGAGGTTAAA and TCAGACCCAGGAATTTGAGG produce 235-bp in the wild type allele and 324-bp fragment in the floxed allele.

*The Wnt1-Cre2* mouse strain was purchased from The Jackson Laboratory (strain #022137) and it was used for specific deletion of the *Meis1* ^fl/fl^ gene in 4V ChP (referred to as *Meis1^KO^* strain in this article). Genotyping primers GCATTTCTGGGGATTGCTTA and CCCGGCAAAACAGGTAGTTA amplify 241-bp Cre fragment.

The reporter line mT/mG was purchased from The Jackson Laboratory (strain #007676).

Mouse strain *Wnt5a^tm1.1Krvl/J^* (referred to *Wnt5a^flox/flox^* in this article) (Ryu et al., 2013) was purchased from Jackson laboratories; *Foxj1^tm1.1(cre/ERT2/GFP)Htg^* (referred to *Foxj1-creERT2* in this article) (Muthusamy et al., 2014) and *Gt(ROSA)26Sor^tm14(CAG-tdTomato)Hze^* (referred to as tdTomato in this article)(Madisen et al., 2010) were shared with the Karolinska Institutet, Sweden through a collaboration agreement. All mice strains were housed, bred and treated in Czech Centre for Phenogenomics (Institute of Molecular Genetics, CAS) in accordance with protocols approved by the animal work committee of the Institute of Molecular Genetics, CAS and Central Commission for Animal Welfare of Ministry of Agriculture Czech Republic (PP-90-2015, PP-64-2018). Induction of conditional knock-out or the tdTomato reporter was induced by single dose of tamoxifen (Sigma) intra-peritoneal injection of pregnant female mice at a concentration of 4.5 mg of tamoxifen dissolved in sterile sunflower oil per 20g weight of mouse.

Overexpression of Wnt5a (*Wnt5a^OE^*) was induced in embryos carrying both an inducible Wnt5a transgene and a Rosa26rtTA driver as described earlier (van Amerongen et al., 2012, strains available via Jackson labs: FVB/N-Tg(tetO-Wnt5a) 17Rva/J, stock number 022938; B6.Cg-Gt(ROSA)26Sortm1(rtTA*M2)Jae/J, stock number 006965). *Wnt5a^OE^* was induced by administrating doxycycline to pregnant mice from E10.5 onwards through dissolving doxycycline in the drinking water (1-2 mg/ml) ad libitum. *Ror2; Vangl2* were obtain from Dr. Yingzi Yang (Gao et al., 2011). Mice were crossed as double heterozygotes (*Ror2^+/-^; Vangl2^+/-^*) due to homozygote lethality. All mice were used according to the rules and regulations of the local ethical committee (Stockholm Norra Djurförsöksetisks Nämd: N273/11, N326/12, N158/15) and (Animal Welfare Committee of the University of Amsterdam).

Reagents used are listed in the Supplementary table 1.

### *In utero* injection

All AAV injection *in utero* experiments were performed under protocols approved by the IACUC of Boston Children’s Hospital (BCH). Timed pregnant CD-1 dams were obtained from Charles River Laboratories. E13.5 CD-1 dams were anaesthetized with isoflurane inhalation and the uterine horns were exposed. 1μl of AAV-GFP or AAV-*Meis1* containing 1∼5X10^12^ gc/ml was injected into a lateral ventricle using a pulled glass pipette (Drummond Scientific Company). Analgesic medication (Meloxicam 5mg/kg) was injected subcutaneously following surgery. E16.5 embryos were harvested and drop fixed in 4% PFA before tissue processing or snap freezing for qPCR analysis.

The murine *Meis1* gene was obtained from the MEIS1A-MIY (Addgene) by EcoRI digestion and ligated into a AAV-CMV-IRES-hrGFP (Agilent) vector digested with EcoRI. AAV5-GFP and AAV5-*Meis1* were produced by the viral core at Boston Children’s Hospital (BCH).

Reagents used are listed in the Supplementary table 1. Constructs used are listed in the Supplementary table 2.

### ChIP-sequencing assays

ChIP-seq protocol was adapted from (Laurent et al., 2015). Briefly, 4V ChP from E18.5 embryos were dissected into cold HBSS. Tissues were crosslinked in 1% formaldehyde at room temperature for 10 minutes while rotating. Glycine (Sigma) was added to a final concentration of 125mM and incubated at room temperature for 5 minutes. Tissues were centrifuged at 4°C at 2000xg and washed twice with 1x PBS with protease inhibitors (Thermo Fisher Scientific). Tissues were snap frozen in liquid nitrogen and stored at −80°C.

Frozen tissues were resuspended in lysis buffer with protease inhibitors (50mM Tris pH 8.1, 10mM EDTA pH 8.0, 1% SDS). Tissues were homogenized using an insulin syringe and then fragmented with an ultrasonic sonicator (Qsonica). After sonication, dilution buffer (0.01% SDS, 1.1% Triton X-100, 1.2mM EDTA, 16.7mM Tris-HCl pH 8.0 and 167mM NaCl) with protease inhibitors were added to give a final SDS concentration of 0.1%.

Chromatin samples were pre-cleared with Protein A beads (Thermo Fisher Scientific) under rotation at 4°C for 1hour prior to incubation overnight with anti-Meis1 and anti-IgG antibody overnight (15 ug of antibody per sample). Protein A beads were added to precipitate antibody complexes and rotated for 1hour at 4°C and then washed with low salt buffer (0.1% SDS, 1% Triton X-100, 2mM EDTA, 20mM Tris-HCl pH 8.0, 150mM NaCl), high salt buffer (0.1% SDS, 1% Triton X-100, 2mM EDTA, 20mM Tris-HCl pH 8.0, 500mM NaCl), LiCl buffer (0.25M LiCl, 1% IGEPAL-CA630, 1% deoxycholic acid, 1mM EDTA, 10mM Tris pH 8.0), and TE buffer (10mM Tris-HCl, 1mM EDTA). Antibody complexes were eluted with 1% SDS in 100mM NaHCO_3,_ and crosslinks were reversed with NaCl and proteinase K (NEB). DNA was recovered by phenol-chloroform extraction and ethanol precipitation and quantified with Qubit (Invitrogen).

ChIP-seq libraries were prepared using NEBNext DNA library preparation reagents (E6000) as described in the Illumina Multiplex ChIP-seq DNA sample Prep Kit. Libraries were indexed using a single indexed PCR primer. Libraries were quantified by Qubit (Invitrogen) and sequenced using a HiSeq 2500 (Illumina) to generate 50 bp single-end reads.

Reagents used are listed in Supplementary table 1.

### ChIP-Sequencing analysis

ChIP-Seq samples were sequenced in single-end on the Illumina HiSeq2500 platform. Raw reads were mapped to mouse reference genomes (mm9) that were downloaded from the UCSC website (www.genome.ucsc.edu). Bowtie (v1.0.0) (Langmead et al., 2009) was used for alignment with “–m 1” and other default parameters. The “-m 1” allows reads uniquely aligned to the genome. MACS (v2.0.10) (Zhang et al., 2008) was used to identify the enriched regions with default parameters. These enriched regions were annotated with an R package, ChIPpeakAnno (v3.6.5) (Zhu et al., 2010) in Bioconductor. The promoters were defined as 2kb from up-to down-stream of transcriptional start sites (TSS).

Reagents used are listed in the Supplementary table 1.

### ChIP-qPCR

ChIP-qPCR was performed on immunoprecipitated DNA after amplification with NEB Next DNA library preparation. qPCR was performed with SYBR green (Roche) for detection on a LightCycler 480 system (Roche) according to the manufacturer’s instructions. The results were calculated as relative fold enrichment over the input.

Reagents used are listed in the Supplementary table 1. Primers are listed in the Supplementary table 5.

### Choroid plexus epithelial cell primary culture

ChP tissue was collected from E14.5 embryos isolated from sacrificed pregnant CD1 mice and choroid plexus epithelial cells (CPEC) were isolated from LV ChP and 4V ChP. During isolation, extracted tissue was kept at room temperature (RT) in HBSS solution (Sigma). After isolation, extracted tissue was briefly centrifuged (200g, 10s at RT). Following aspiration of supernatant, 500µl of 2 mg/ml solution of Pronase (Sigma) was added to the extracted tissue and incubated for 5 minutes at 37°C. The solution was then transferred to DMEM/F-12 medium containing 10% FBS (Thermo Fisher Scientific) and centrifuged (300g, 2min at RT). Tissue was transferred to complete culture medium consisting of DMEM/F-12 supplemented with 10% FBS, 10ng/ml EGF (Invitrogen), 20 µM cytosine arabinoside (Sigma) 50U/ml penicillin, and 50U/ml streptomycin. Cells were mechanically dissociated through a 21-gauge needle using 6-8 times forced passages, followed by gentle repeated resuspension with a 200µl pipette. Finally, cells were seeded onto Transwell-0,4μm thick clear filter inserts (Sigma). Inserts were pre-coated on their upper side with laminin (Sigma) as described by the manufacturer. To achieve higher purity of epithelial cells, an adhering-off method was applied to reduce fibroblast contamination. After the initial seeding, supernatant containing unadhered cells was transferred to new laminin coated well thus removing from culture, fibroblasts characterized by a higher adherence affinity.

In order to produce CM, CPEC primary cultures were maintained in complete culture medium. CM was collected every 48 hours up to 10 days after seeding. Supernatant was subjected to sequential centrifugation steps of 200g for 5 min (to remove viable cells), 1500g for 10 minutes (to remove death cells) and 6000g for 15 minutes (to remove cell debris).

Reagents used are listed in the Supplementary table 1.

### Fetal tissue section

Ethical approval allowing human fetal tissue acquisition and analysis was provided by the National Research Ethics Service Committee East of England—Cambridge Central, UK (ethics number 96/085).

### Cell culture and transfection

HEK293T cells were seeded in complete DMEM medium containing 10% FBS, 2mM L-glutamine, 50U/ml penicillin, and 50U/ml streptomycin (Thermo Fisher Scientific) on 10cm dishes 24 hours prior transfection. The cells were transfected with a total of 5µg of DNA at ∼40% confluency in DMEM medium only. The transfection reaction mixture was prepared using OptiMEM (Thermo Fisher Scientific) and Lipofectamine 2000 (Thermo Fisher Scientific), with a ratio of 1µg DNA: 2µl Lipofectamine 2000, followed by incubation with cells for 4-6 hours. Afterwards the transfection medium was exchanged for the complete medium.

Reagents used are listed in the Supplementary table 1.

### CRISPR/Cas9 generation of WNT5A-only HEK293 T-REx cells

Plasmid encoding guide RNA targeting the human WNT5A gene was used. gRNA sequence was cloned into plasmids pSpCas9(BB)-2A-GFP (PX458) (Addgene plasmid, 41815) or pU6-(BbsI) CBh-Cas9-T2A-mCherry (Addgene plasmid, 64324). T-REx-293 cells (Invitrogen) were cultured according to the manufacturer’s instructions and transfected by Lipofectamine® 2000 DNA Transfection Reagent (Thermo Fisher Scientific) with plasmids encoding guide RNAs targeting human WNT5A gene. Next day, transfected cells were single cell sorted and grown as single colonies. Selection of WNT5A knock-out (*^KO^*) clones was done by PCR. Genomic DNA was isolated by DirectPCR Lysis Reagent (Viagen Biotech) and then

the fragment of genomic DNA was amplified by PCR using DreamTaq DNA Polymerase (Thermo Fisher Scientific). Then, the PCR product was cut by TaqI (Thermo Fisher Scientific). WNT5A ablation efficiency was assessed by Western Blot analysis using antibodies: WNT5A antibody (R&D). All cell lines were verified and sequenced.

Reagents used are listed in the Supplementary table 1. Constructs used are listed in the Supplementary table 2. Primers are listed in the Supplementary table 5.

### Western blotting

Samples were subjected to SDS-PAGE, electrotransferred onto a Hybond-P membrane (GE Healthcare), immunodetected using appropriate primary and secondary antibodies and visualized by ECL (Millipore) or Supersignal Femto solution (Thermo Fisher). Signal intensities were calculated using ImageJ. Briefly, the area of the peak intensity for the protein of interest was divided by the corresponding values of peak intensity obtained for the control protein.

Reagents used are listed in the Supplementary table 1. Antibodies used are listed in the Supplementary table 3.

### Micro CT

Embryos were fixed for 1 week in 4% PFA and stained with Lugol’s Iodine solution for 2 weeks or longer. Stock solution (10g KI and 5g I2 in 100ml H2O) was diluted to 25% working solution in H2O. Stained specimens were removed from contrast agent, rinsed with PBS and embedded in 2.5% low gelling temperature agarose dissolved in water. Scan was performed on SkyScan 1272 high-resolution microCT (Bruker, Belgium), with voxel size set up for 3µm.

### Immunofluorescence and EdU staining

For mouse embryo immunofluorescent analysis and *in situ* hybridization, WT CD1 mice, *Wnt5a*^-/-^ mice were dissected and isolated embryos were transferred into ice-cold PBS, fixed in 4% paraformaldehyde (PFA, Millipore) in PBS for several hours followed by several washes in ice-cold PBS and finally cryoprotected by sequential incubation in 15% and then 30% sucrose solutions. Embryos were next frozen in optimum cutting temperature (OCT) compound (Sakura FineTek) on dry ice. Serial 14µm coronal sections were used for immunofluorescence analysis. Human fetal tissue for cryosectioning was immersion-fixed overnight in 4% PFA at 4°C, then cryoprotected in sucrose before embedding in the OCT compound and then 14 µm sections were cut using a Leica cryostat. Human fetal tissue samples were then processed using an identical immunofluorescence protocol as for the mouse samples.

For immunofluorescent analysis, all the sections underwent antigen retrieval by direct boiling for 1 minute at 550W in the microwave followed by 10 minutes incubation at 85°C in water bath, using antigen retrieval solution (DAKO). Sections were washed in PBT (PBS with 0.5% Tween-20) and blocked in PBTA (PBS, 5% donkey serum, 0.3% Triton X-100, 1% BSA). Samples were incubated overnight at 4°C with primary antibodies diluted in PBTA. Following washes in PBT, samples were incubated with corresponding Alexa Fluor secondary antibodies (Invitrogen, Abcam) for 1h at RT, followed by 5 minutes incubation at RT with DAPI (Thermo Fisher). Finally, samples were mounted in DAKO mounting solution (DAKO).

EdU (Life Technologies) was injected 72 hours before the embryos were harvested at a concentration of 65 mg/g. Cells with incorporated EdU were visualized using a Click-iT EdU Alexa Fluor 555 Imaging Kit (Life Technologies).

For immunocytofluorescent analysis of ChP primary culture, cells grown on laminin coated cover slips were first washed several times in ice-cold PBS, followed by fixation for 15 minutes in ice-cold 4% PFA. Later, cells were washed several times in PBT, blocked with PBTA for 30 minutes and incubated overnight at 4°C with primary antibodies. Following repeated washing in PBT, cells were incubated for 1 hour at RT with appropriate secondary antibodies, DAPI for 5 minutes and mounted in DAKO mounting medium (DAKO).

Reagents used are listed in the Supplementary table 1. Antibodies used are listed in the Supplementary table 3.

### *In situ* analysis

*In situ* analysis of the gene expression was performed on 14-µm cryosections of embryos at various stages of embryonic development isolated from CD1 mice. After isolation, embryos were immediately transferred and kept in fresh 4% PFA for 2 hours, washed briefly in ice cold PBS, incubated for 6 hours in 30% sucrose solution at 4°C, and frozen at −80°C. Transcripts were detected using an adapted protocol for the RNAscope 2.0 assay for fixed frozen tissue (Advanced Cell Diagnostics). Staining was performed using the RNAscope Fluorescent Multiplex Reagent Kit (Advanced Cell Diagnostics).

Indicated *in situ* images were adopted from Allen Institute for Brain Science: Allen Developing Mouse Brain Atlas (Lein et al., 2006) (Available from: http://developingmouse.brain-map.org) or from eurexpress.org (Diez-Roux et al., 2011) (Available from: http://www.eurexpress.org/ee/).

Reagents used are listed in the Supplementary table 1. Probes used are listed in the Supplementary table 4.

### Real time qPCR

RNA was isolated using from 3 different litters of WT *CD1* embryos collected at different developmental stages using RNAeasy kit (Qiagen). Samples were treated with DNase (Qiagen) to prevent contamination with genomic DNA. The specificity of primers was determined by BLAST run of the primer sequences. Annealing temperature was 57°C for all used primers.

qPCR reactions were performed once for every gene / sample in triplicate. PCR was done according to the manufacturer’s protocol using Lightcycler 480 SYBR Green 1 Master Mix (Roche). The following thermo cycling program parameters were used for qPCR analysis: Incubation step at 95°C for 5min, then 45 cycles 95°C for 10 seconds, 57°C for 10 seconds and 72°C for 10 seconds. qPCR analysis was carried out using LightCycler© 480 Instrument II (Roche).

ΔCp values were calculated in every sample for each gene of interest with *β-Actin* as the reporter gene. Relative change of expression level for analyzed gene (ΔCp) was performed by subtraction of the gene expression levels in the LV ChP or 4VChP from the gene expression level of housekeeping gene (*β-actin*). Next, the ratio of the gene expression level between *β-actin* and gene of interest in either 4V ChP or LV ChP was calculated using the following formula: 2^-^ΔCp.^

Reagents used are listed in the Supplementary table 1. Primers are listed in the Supplementary table 5.

### Statistics

Gene expression data - Replicates represent independent experiments. Data in Fig. 3F, Fig. 4C, Fig. 5A and Supplementary Fig. 4D are represented in columns showing the mean with standard deviation (s.d.). Significance was measured using a paired two-tailed Student’s t-test with unequal sample variance. Biological replicates per condition are indicated in the corresponding graphs. Data in Fig. 5G are represented as columns showing the standard error of the mean (s.e.m).

Confocal images *-* Data in Fig. 1G-I, Fig. 1M-O, Fig. 2E-G, Fig. 2J-K, Fig. 6C, Fig. 6F-G, Fig. 6J-K, Supplementary Fig. 1E-F, Supplementary Fig. 1H and Supplementary. Fig. 2J-K are expressed as columns showing the mean with s.d. Significance was measured using unpaired or paired two-tailed Student’s t-test with unequal sample variance. Biological replicates used per condition are indicated in the corresponding graphs.

Immunoblot images - Images used for quantitative analysis reported Supplementary Fig. 4A, C and Supplementary Fig. 5D are expressed as columns showing the mean with s.d. Significance was measured using unpaired two-tailed Student’s t-test with unequal sample variance. Biological replicates per condition are indicated in the corresponding graph.

Except for Fig.6 J-K, Supplementary Fig. 1I-J, Supplementary Fig. 2C and Supplementary Fig. 2G all the displayed immunostaining images and western blots are representative of at least 3 independent experiments.

## ACKNOWLEDGEMENTS

We thank members of Bryja and Lehtinen labs for their help and suggestions, Dr. Marcela Buchtová and Mgr. Marie Šulcová (Masaryk University) for their valuable assistance with RNAscope hybridization, Dr. Sean Li (Boston Children’s Hospital) for sharing *Wnt5a* knockout mice with the Lehtinen lab, and Chen Wu at the BCH Viral Core. We are also thankful to Nikodém Zezula (Masaryk University) for his assistance with the figure graphic design and Mgr. Monika Novákova (Masaryk University) for help with the animal work. We thank MEYS CR for support to the following core facilities: CELLIM of CEITEC supported by the Czech-BioImaging large RI project (LM2018129 funded by MEYS CR), Czech Centre for Phenogenomics (LM2015040), Higher quality and capacity of transgenic model breeding (by MEYS and ERDF, OP RDI CZ.1.05/2.1.00/19.0395), Czech Centre for Phenogenomics: developing towards translation research (by MEYS and ESIF, OP RDE CZ.02.1.01/0.0/0.0/16_013/0001789), BCH viral core for AAV production. The collaboration between Masaryk University and Karolinska Institutet (KI-MU program), was co-financed by the European Social Fund and the state budget of the Czech Republic (CZ.1.07/2.3.00/20.0180). Funding to the VB lab was obtained from Neuron Fund for Support of Science (23/2016), and Czech Science Foundation (GA17-16680S). Work in the EA lab was supported by the Swedish Research Council (VR projects: DBRM, 2011-3116, 2011-3318 and 2016-01526), Swedish Foundation for Strategic Research (SRL program and SLA SB16-0065), European Commission (NeuroStemcellRepair), Karolinska Institutet (SFO Strat Regen, Senior grant 2018), Hjärnfonden (FO2015:0202 and FO2017-0059) and Cancerfonden (CAN 2016/572) foundations. KK was supported by Masaryk University (MUNI/E/0965/2016). RvA acknowledges funding support from the University of Amsterdam (MacGillavry fellowship), KWF Kankerbestrijding (Dutch Cancer Society, career development award ANW 2013-6057) and NWO (Netherlands Science Foundation, VIDI 864.13.002).Work in the OM lab is supported by the Czech Science Foundation (18-00514S). ZK acknowledges funding support from GACR 18-20759S. RA Barker is supported by an NIHR funded Biomedical Research Centre at Cambridge University Hospital and the WT-MRC Cambridge Stem Cell Institute. The Lehtinen laboratory was supported by NIH R01 NS088566 (MKL), the New York Stem Cell Foundation (MKL); and BCH IDDRC 1U54HD090255. MK Lehtinen is a New York Stem Cell Foundation – Robertson Investigator.

## AUTHOR CONTRIBUTIONS

KK designed and performed most of the experiments, analyzed data, prepared all figures, and wrote the manuscript; AJ and MPL prepared the AAV constructs, performed *in utero* injections, qPCR and immunoblotting; MPL; ML and BL performed chromatin immunoprecipitation and analyzed the data with FW; JP, MP, IU and RS produced *Wnt5a^cKO^* strain; OM and ZK produced the *Meis1^KO^* strain; RvA provided the *Wnt5a^OE^* embryos, DG and EA provided the *Wnt5a^KO^* and *Vangl2^KO^* embryos, PK performed immunostaining analysis; RLG and RAB contributed the human fetal samples; VB and ML designed experiments, supervised the work, analyzed data and wrote the manuscript.

---

**Supplementary Table 1.**

| Reagent | Manufacturer | Cat. no. |
| --- | --- | --- |
| Amersham Hybond P 0.45 PVDF blotting membrane | GE healthcare | 10600023 |
| Antigen retrieval solution | DAKO | S1699 |
| DAPI (4',6-Diamidino-2-Phenylindole, Dihydrochloride) | Thermo Fisher Scientific | D1306 |
| DirectPCR Lysis Reagent | Viagen Biotech | 301-C |
| DreamTaq DNA Polymerase | Thermo Fisher Scientific | EP0701 |
| EdU (5-ethynyl-2'-deoxyuridine) | Thermo Fisher Scientific | A10044 |
| EdU Alexa Fluor 555 Imaging Kit | Thermo Fisher Scientific | C10338 |
| Fetal Bovine Serum | Thermo Fisher Scientific | 16000044 |
| Flourescence mounting medium | DAKO | S3023 |
| Formaldehyde solution 4%, buffered, pH 6.9 | Millipore | 1004960700 |
| Hanks' Balanced Salt solution | Sigma | H9394-500ML |
| Immobilon ECL Ultra Western HRP Substrate | Millipore | WBULS0500 |
| Laminin | Sigma | L2020 |
| L-Glutamine | Thermo Fisher Scientific | 21051024 |
| LightCycler® 480 SYBR Green I Master | Roche | 4707516001 |
| Lipofectamine 2000 transfection reagent | Thermo Fisher Scientific | 11668-019 |
| Meis1 TaqMan probe Mm00487664_m1 | Thermo Fisher Scientific | 4331182 |
| Opti-MEM™ I Reduced Serum Medium | Thermo Fisher Scientific | 31985-047 |
| Penicillin-Streptomycin | Thermo Fisher Scientific | 15140122 |
| Pronase | Sigma | 10165921001 |
| RNAscope Fluorescent Multiplex Reagent | Advanced Cell Diagnostics | 320850 |
| RNase-Free DNase Set | Qiagen | 79254 |
| RNeasy Mini Kit (50) | Qiagen | 74104 |
| SuperSignal™ West Femto Maximum Sensitivity Substrate | Thermo Fisher Scientific | 34094 |
| Tamoxifen | Sigma | T5648-1G |
| Taq1 restriction enzyme | Thermo Fisher Scientific | ER0671 |
| Wnt5a TaqMan probe Mm00437347_m1 | Thermo Fisher Scientific | 4331182 |
| Tissue-Tek® O.C.T. Compound, Sakura® Finetek | Sakura Finetek | 25608-930 |
| Transwell-0,4µm thick clear filter inserts | Sigma | CLS3397 |
| T-REx™-293 Cell Line | Thermo Fisher Scientific | R71007 |

**Supplementary Table 2.**
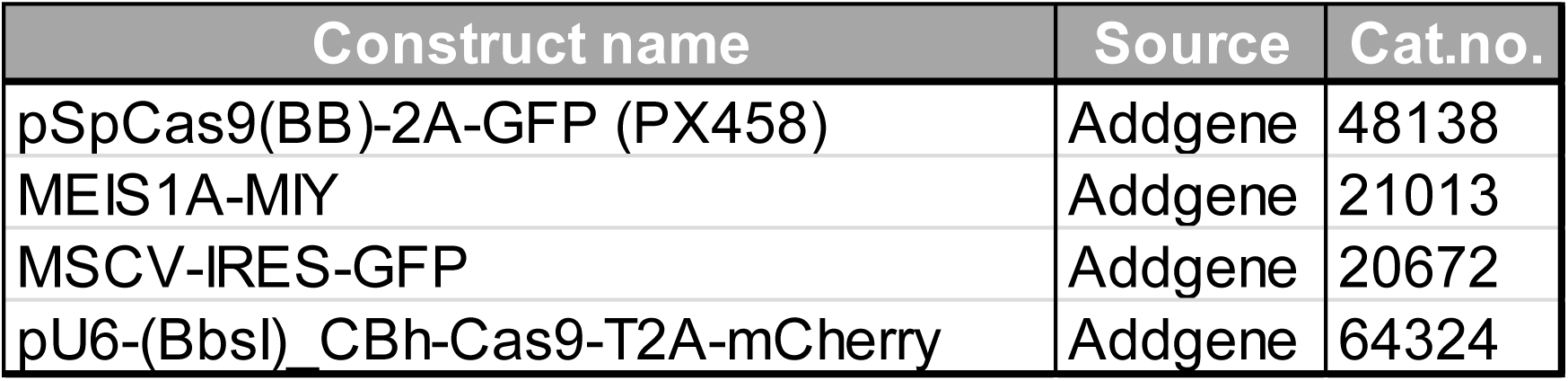

**Supplementary Table 3.**

| Name | Manufacturer | Cat.no. | Species | Application | Dilution |
| --- | --- | --- | --- | --- | --- |
| 555 Alexa Fluor Mouse | Abcam | ab150158 | Goat | IF | 1:750 |
| 488 Alexa Fluor Rabbit | Thermo Fisher Scientific | A21206 | Donkey | IF | 1:750 |
| 680 Alexa Fluor Mouse | Thermo Fisher Scientific | A32788 | Donkey | IF | 1:750 |
| AQP-1 | Santa Cruz | sc-55466 | Mouse | IF | 1:500 |
| Caspase-3 | Cell Signaling | cs-9664 | Rabbit | IF | 1:500 |
| Claudin-1 | Invitrogen | 51-9000 | Rabbit | IF | 1:1000 |
| Dvl2 | Cell Signaling | cs-3216 | Rabbit | WB | 1:500 |
| Dvl3 | Santa Cruz | sc-8027 | Mouse | WB | 1:500 |
| FLAG | Sigma | F7425 | Rabbit | IF | 1:1000 |
| GAPDH | Cell Signaling | 97166S | Rabbit | WB | 1:1000 |
| KI67 | Invitrogen | 14-5698-80 | Rat | IF | 1:250 |
| total LRP6 | Cell Signaling | cs-2560 | Rabbit | WB | 1:500 |
| p-LRP6 | Cell Signaling | cs-3395S | Rabbit | WB | 1:500 |
| Meis1 | Atlas | HPA056000 | Rabbit | IF (Human) | 1:250 |
| Meis1 | Abcam | ab19867 | Rabbit | WB, IF (Mouse) | 1:250 |
| p-LRP6 | Cell Signaling | cs-2568 | Rabbit | WB | 1:500 |
| Ror1 | kind gift from Henry Ho |  | Rabbit | WB | 1:1000 |
| Ror2 | Sigma | HPA021868 | Rabbit | WB | 1:1000 |
| Scribble | GeneTex | GTX107692S | Rabbit | IF | 1:250 |
| Vangl2 | Millipore | MABN750 | Rat | WB, IF | 1:200 |
| WNT5A | R&D | MAB645 | Rat | WB, IF | 1:250 |
| β-Actin | Cell Signaling | 4970 | Rabbit | WB | 1:1000 |

**Supplementary Table 4.**

| Gene | Species | Manufacturer | Cat.no. |
| --- | --- | --- | --- |
| *Axin2* | mouse | ACDBio | 400331-C3 |
| *Foxj1* | mouse | ACDBio | 317091 |
| *Lef1* | mouse | ACDBio | 441861 |
| *Meis1* | mouse | ACDBio | 436361-C2 |
| *Meis2* | mouse | ACDBio | 43678 |
| *Ror1* | mouse | ACDBio | 445301 |
| *Ror2* | mouse | ACDBio | 430041 |
| *Vangl2* | mouse | ACDBio | 316781 |
| *Wnt5a* | mouse | ACDBio | 316791-C3 |

**Supplementary Table 5.**
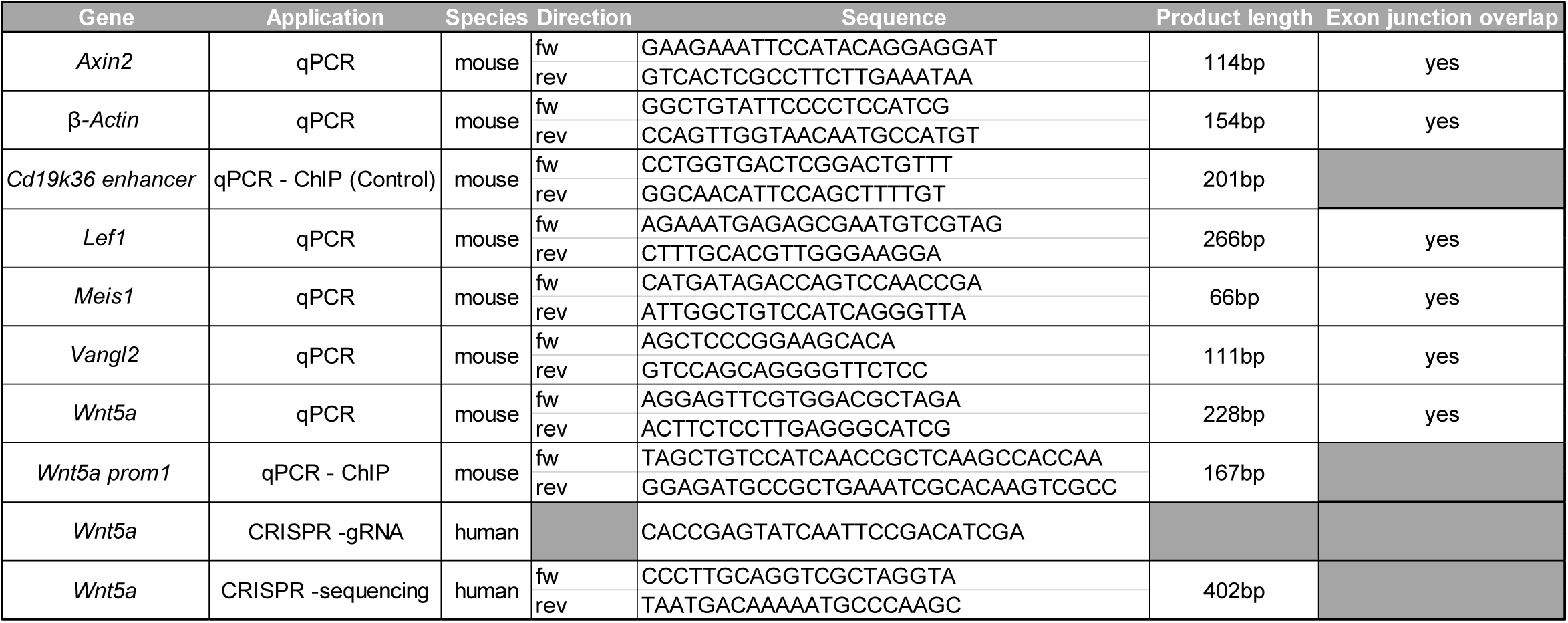

